# In-depth single-cell analysis of translation-competent HIV-1 reservoirs identifies cellular sources of plasma viremia

**DOI:** 10.1101/2021.02.15.431218

**Authors:** Basiel Cole, Laurens Lambrechts, Pierre Gantner, Ytse Noppe, Noah Bonine, Wojciech Witkowski, Lennie Chen, Sarah Palmer, James I. Mullins, Nicolas Chomont, Marion Pardons, Linos Vandekerckhove

## Abstract

Clonal expansion of HIV-infected cells contributes to the long-term persistence of the HIV reservoir in ART-suppressed individuals. However, the contribution to plasma viremia from cell clones that harbor inducible proviruses is poorly understood. Here, we describe a single-cell approach to simultaneously sequence the TCR, integration sites and proviral genomes from translation-competent reservoir cells, called STIP-Seq. By applying this approach to blood samples from eight participants, we showed that the translation-competent reservoir mainly consists of proviruses with short deletions at the 5’-end of the genome, often involving the major splice donor site. TCR and integration site sequencing revealed that antigen-responsive cells can harbor inducible proviruses integrated into cancer-related genes. Furthermore, we found several matches between proviruses retrieved with STIP-Seq and plasma viruses obtained during ART and upon treatment interruption, showing that STIP-Seq can capture clones that are responsible for low-level viremia or viral rebound.

## Introduction

HIV-1 infection remains incurable due to the establishment of a persistent viral reservoir, which is unaffected by antiretroviral therapy (ART)^1–4^. This reservoir mainly consists of long-lived memory CD4 T cells harboring latent, replication-competent proviruses, capable of refueling viremia upon treatment interruption (TI)^5–7^. The viral reservoir is remarkably stable, with an estimated half-life of ~44-48 months, suggesting that at least 70 years of continuous ART would be required to eliminate it completely^5,8,9^. Long-term maintenance of the reservoir can in part be explained by clonal expansion of HIV-infected cells, which is thought to be driven by three non-mutually exclusive forces: homeostatic proliferation^10–15^, antigenic stimulation^16–18^, and integration site-driven proliferation^19–22^. Identifying the cellular sources of viral rebound and the mechanisms that ensure their persistence during ART is needed to develop targeted strategies to eradicate or control HIV^23–26^.

Several sequencing-based assays have been developed to study the HIV reservoir, each focusing on different aspects of the infected cells and the proviruses within^27^. Near full-length (NFL) provirus sequencing enables the identification of genome-intact and potentially replication-competent proviruses^28–31^. Integration site analysis (ISA) pinpoints the chromosomal location of proviruses and is frequently used as a marker to study clonal expansion of infected cells^19,20,23,32^. More recently, NFL provirus sequencing and ISA were combined into a single assay, allowing the study of the relationship between proviral integration site (IS) and genome structure^33,34^. However, because these assays are usually performed on bulk CD4 T cell DNA, they mainly identify defective proviruses, as it has been estimated that only 2-5% of the total proviruses are genome-intact^28,35–37^. As such, they do not focus on proviruses that could lead to viral rebound upon TI. On the contrary, viral outgrowth assays (VOA) combined with NFL viral genome sequencing enable the characterization of replication-competent proviruses^3,8,38,39^. However, the IS of the provirus as well as the phenotype and TCR sequence of the infected cell cannot be determined with this assay.

Alternative assays have been developed to characterize and quantify infected cells harboring transcription-competent^22,40^ or translation-competent^17,40–44^ proviruses, therefore enriching for proviruses with a higher probability of contributing to viral rebound^45^. These assays use a potent stimulant to reactivate proviruses from latency, inducing transcription of viral genes and production of viral proteins. Infected cells can then be identified and isolated by fluorescence-activated cell sorting (FACS). This allowed for the characterization of NFL proviral genome structure^22,43^, TCR sequences^17,43^ and IS^22^ from cells harboring an inducible provirus. However, none of these methodologies capture all three layers of information simultaneously.

Here, we present a novel method, called HIV STIP-Seq: <u>S</u>imultaneous <u>T</u>CR, <u>I</u>ntegration site and <u>P</u>rovirus sequencing. STIP-Seq enables sequencing of the proviral genome and matched IS of translation-competent proviruses, as well as phenotypic characterization and TCR sequencing of the host cell. We used this approach to characterize infected cells that harbor inducible proviruses from 8 individuals on suppressive ART. Furthermore, 3 out of 8 participants underwent an ATI, which allowed us to investigate the contribution of the translation-competent reservoir to residual viremia and viral rebound.

## Results

### STIP-Seq

STIP-Seq is a derivative of the HIV-Flow assay^41^, with the addition of downstream whole genome amplification (WGA) and sequencing of the provirus, IS and TCR. Since WGA by multiple displacement amplification (MDA) is not compatible with cross-linking fixatives such as paraformaldehyde, we used methanol for simultaneous fixation and permeabilization, permitting efficient amplification of the cellular genome. Using a dilution series of J1.1 cells in the parental Jurkat cell line, we showed good linearity of the frequency of p24+ cells assessed by the methanol-based HIV-Flow assay, down to ~3 p24+ cells/million cells (R^2^ =0.99, Supplementary Fig. 1a, b). In addition, methanol fixation did not have a significant impact on the frequency of p24+ cells (p=0.84, Supplementary Fig. 1c).

Following methanol-based HIV-Flow, p24+ cells were sorted into individual wells of a 96-well plate (Fig. 1a, b). Single-cell whole genome amplification by MDA was used to amplify the DNA of single sorted p24+ cells, including the provirus integrated within. Amplified genomes were subjected to ISA by Integration Site Loop Amplification (ISLA) and NFL proviral sequencing using either a 5− or 2-amplicon PCR approach (Fig. 1a, Supplementary Fig. 2). In addition, the TCRβ chain of the host cell was sequenced as described^17^, and index sorting was used for *post hoc* determination of the memory phenotype of p24+ cells (Fig. 1a, Supplementary Fig. 2).

**Figure 1:**
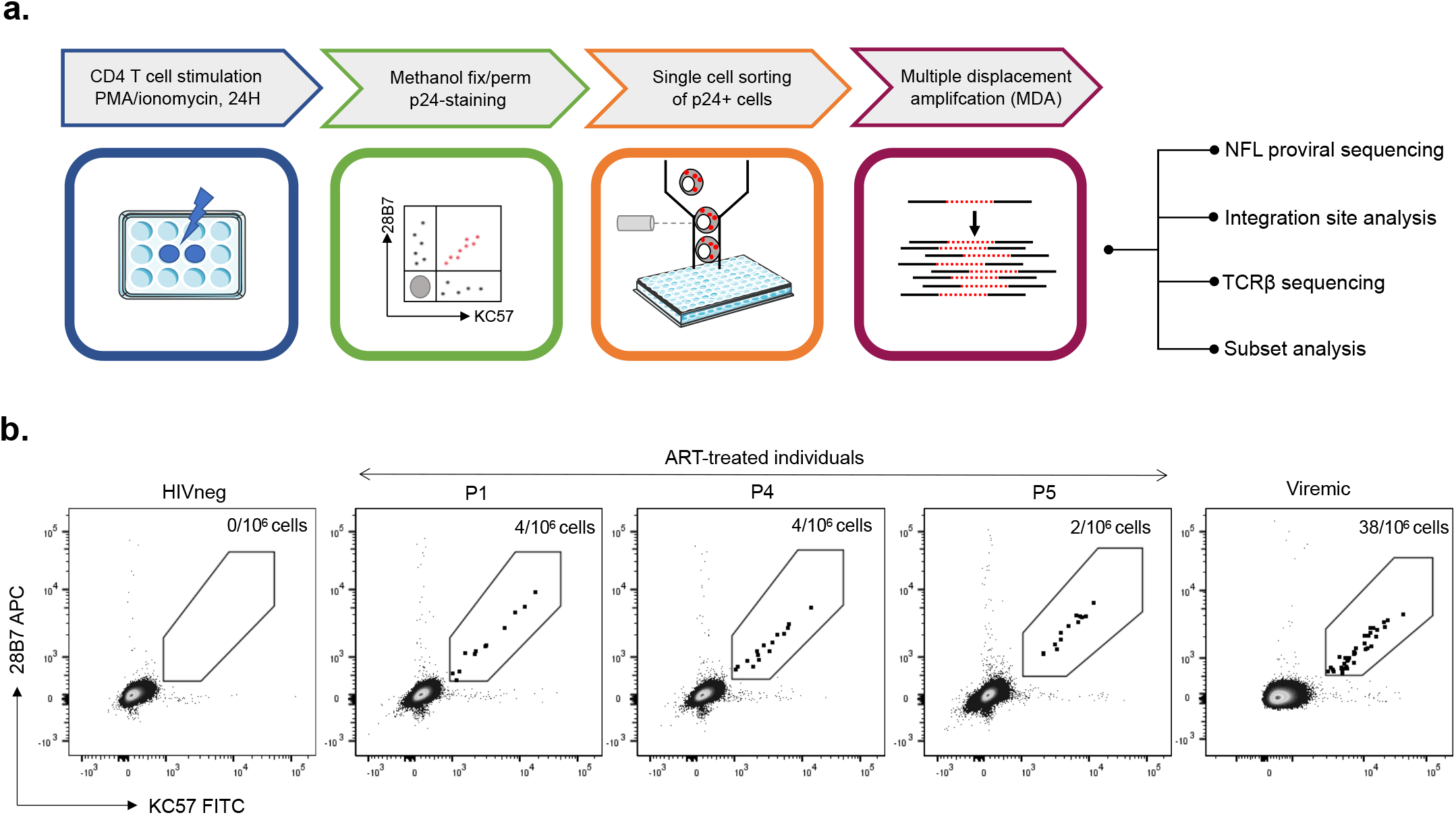
STIP-Seq enables isolation and characterization of p24-producing cells after PMA/ionomycin stimulation. (**a**) Overview of the STIP-Seq assay. CD4 T cells are stimulated for 24 h with PMA (162nM) / ionomycin (1ug/mL). Cells are fixed and permeabilized with methanol, and p24-producing cells are identified using a combination of 2 antibodies (KC57 and 28B7) targeting the p24 protein. p24+ cells are single-cell sorted by flow cytometry. DNA from p24+ cells is amplified by multiple displacement amplification, before performing near full-length (NFL) proviral genome sequencing, integration site analysis, TCR sequencing, and *post-hoc* determination of the CD4 T cell memory phenotype. NFL = near full-length. (**b**) Representative FACS dot plots showing the KC57-FITC/28B7-APC co-staining on CD4 T cells from 1 HIV non-infected control, 1 viremic and 3 ART-treated individuals.

### STIP-Seq enables deep characterization of the translation-competent HIV-1 reservoir in ART-suppressed individuals

To investigate the characteristics of p24-producing cells and their associated proviruses, we performed STIP-Seq on single sorted CD4 T cells from 8 ART-suppressed individuals (Supplementary Table 1). Of note, for participant P5, STIP-Seq was performed on 2 samples collected 3 years apart. A total of 158 p24+ cells and 156 IS were retrieved. A large proportion of these stemmed from clonally expanded infected cells (74%, 116/156), defined by recurrent identical IS, confirming the often clonal nature of the translation-competent reservoir^17^.

NFL proviral genome sequencing yielded a total of 40 distinct genomes with complete coverage, which fell within one of three categories: genome-intact (12.5%, 5/40), packaging signal (PSI) and/or major splice donor (MSD) defects (85%, 34/40), or large internal deletion (2.5%, 1/40) (Fig. 2a, Supplementary Table 2). The PSI/MSD defective proviruses usually had deletions spanning one or more packaging stem-loops, all of them involving the MSD located within stem-loop 2 (Fig. 2b). Among these, we identified 7 proviruses with deletions covering the binding region of the forward primer from the 5-amplicon NFL PCR (U5-638F), although these deletions could be spanned by the 2-amplicon approach (F581; Fig. 2b, indicated with triangles). Intriguingly, 16 proviruses (40%) had deletions extending into the *p17* gene, removing the start-codon of the Gag polyprotein (Supplementary Fig. 3). This implies the use of an alternative start-codon to enable the translation of the p24 protein^46,47^. Out of 40 distinct NFL sequences analyzed, only 1 had a large internal deletion (1191 bp; Fig. 2a), and none displayed inversions or hypermutations (Fig. 2a). This is in contrast with previously reported NFL data that were generated on bulk CD4 T cell DNA^28–30,35^. To investigate this disparity, we compared NFL genomes obtained by *Full-Length Individual Provirus Sequencing* (FLIPS)^28^ on bulk CD4 T cell DNA with NFL sequences retrieved by STIP-Seq, for two longitudinal samples from participant P5 (Supplementary Fig. 4). This analysis showed that proviruses with large internal deletions and hypermutations were absent in p24+ cells, although highly prevalent in bulk CD4 T cells (1/65 hypermutated, 58/65 deleted, Supplementary Fig. 4). At the second time point, 2/65 proviral genomes recovered from bulk CD4 T cells were intact, whereas none were detected in p24+ cells (0/11) (Supplementary Fig. 4). This suggests that these proviruses were not induced by a single round of PMA/ionomycin stimulation, or they were missed due to the more limited sampling with STIP-Seq.

**Figure 2:**
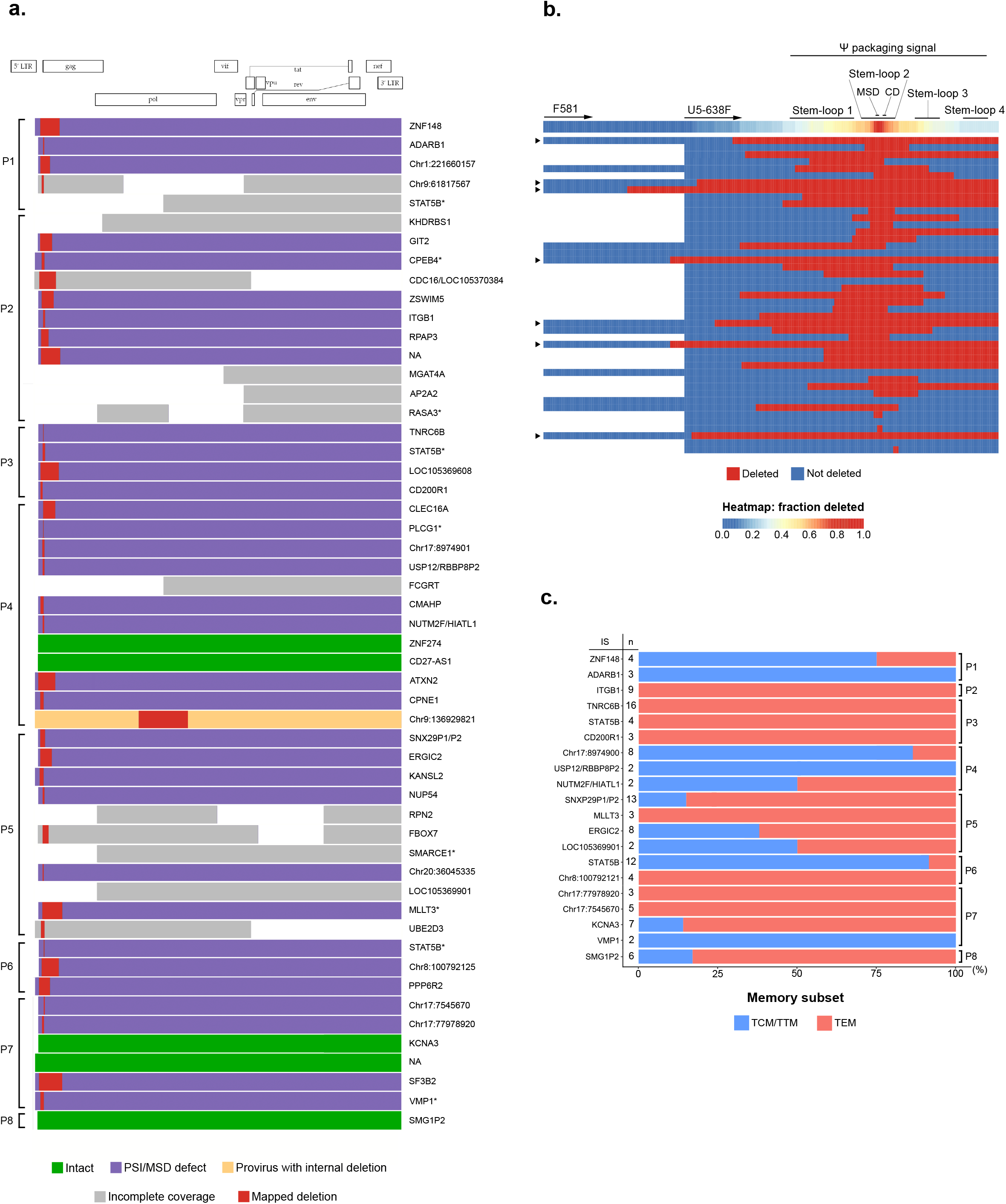
Near full-length proviral sequencing, integration site analysis and subset analysis on p24-producing cells from ART-treated individuals. (**a**) Virogram showing the near full-length proviral genomes recovered from 8 ART-treated individuals. Proviral genomes were reconstructed using a 5-amplicon, 2-amplicon or 4-amplicon PCR-approach. Corresponding integration sites (IS) are indicated at the right-hand side of each proviral genome. Cancer-related genes are indicated with an asterisk. (**b**) Heatmap of the deletions in the 5’ UTR region, including the Ψ packaging signal. The second-round forward primers for the 2-amplicon (F591) and 5-amplicon (U5-638F) NFL PCR-approach are annotated with arrows on the heatmap. Proviruses with a deletion spanning the U5-638F primer are indicated with a triangle at the left-hand side of each provirus. MSD = major splice donor, CD = cryptic donor. (**c**) Memory subset distribution of clonal p24-producing cells. The number of cells within each clone is indicated at the left-hand side of each horizontal bar. TCM = central memory T cell, TTM = transitional memory T cell, TEM = effector memory T cell.

In order to link the chromosomal location of proviruses to their corresponding genome structure, ISA was performed on successfully amplified genomes. A bias towards integration in the reverse orientation with respect to the gene was observed (36/58 in reverse orientation, 12/58 in same orientation, 3/58 in region with gene on either strand, 9/58 in intergenic region) (Supplementary Table 3). Previous studies have shown an enrichment of IS in cancer-associated genes, such as *STAT5B* and *BACH2*, suggesting IS-driven expansion of infected cells^19–21,48,49^. Out of 58 distinct IS, 11 were located within cancer-associated genes (Fig. 2a, Supplementary Table 3, indicated with asterisks). Among those, three different IS in the *STAT5B* gene were identified, two of which could be attributed to clonally expanded cell populations. Of note, all three proviruses were integrated in the opposite orientation with respect to the gene. Interestingly, for participant P4, a cell with an intact provirus integrated in the *ZNF274* gene was retrieved (Fig. 2a). This gene was previously described as located in a dense heterochromatin region and was associated with proviruses in a state of ‘deep latency’^50^.

It was previously shown that p24+ cells mainly display central memory (TCM), transitional memory (TTM) and effector memory (TEM) phenotypes^17,41^. Consistent with this, all but one of the cells identified with STIP-Seq fell within these memory subsets (60/143 TCM/TTM, 82/143 TEM), with a single cell displaying a naïve phenotype (1/143 TN). When restricting the analysis to clones, 9/20 were found in both the TCM/TTM and the TEM subset (Fig. 2c), an observation that was previously reported^17,18^. Of note, proportions of CD4 T cell subsets were only minimally affected by a 24h PMA/ionomycin stimulation and methanol fixation (Supplementary Fig. 5).

In conclusion, these results show that p24+ cells preferentially display a memory phenotype and are enriched in NFL proviral genomes that have deletions at the 5’ end of the genome. Our data suggest that the MSD, located within stem-loop 2, is a particular hotspot for deletion among translation-competent proviruses.

### TCRβ sequencing reveals clonal infected cell populations with predicted responsiveness towards pathogens

Under the hypothesis that infected cell clones with responsiveness towards a pathogen could have arisen due to cognate antigen exposure, we attempted to predict the specificity of p24+ cells based on the CDR3 region of the TCRβ sequence, as described^17^. A total number of 43 distinct TCRβ sequences were retrieved. Importantly, p24+ cells that were previously determined clonal by IS sequencing were also identified as such based on TCRβ sequences. The proportion of HIV-infected cells for which specificity could be predicted was 8/43 (19%) when considering all distinct CDR3 sequences, or 5/19 (26%) when restricting to clonal populations (Fig. 3, Supplementary Table 4). Among all participants, predicted TCR specificities of p24+ cells were confined to CMV, *M. tuberculosis* and influenza, suggesting that infection with or immunization against these pathogens plays a role in the maintenance of the translation-competent reservoir.

**Figure 3:**
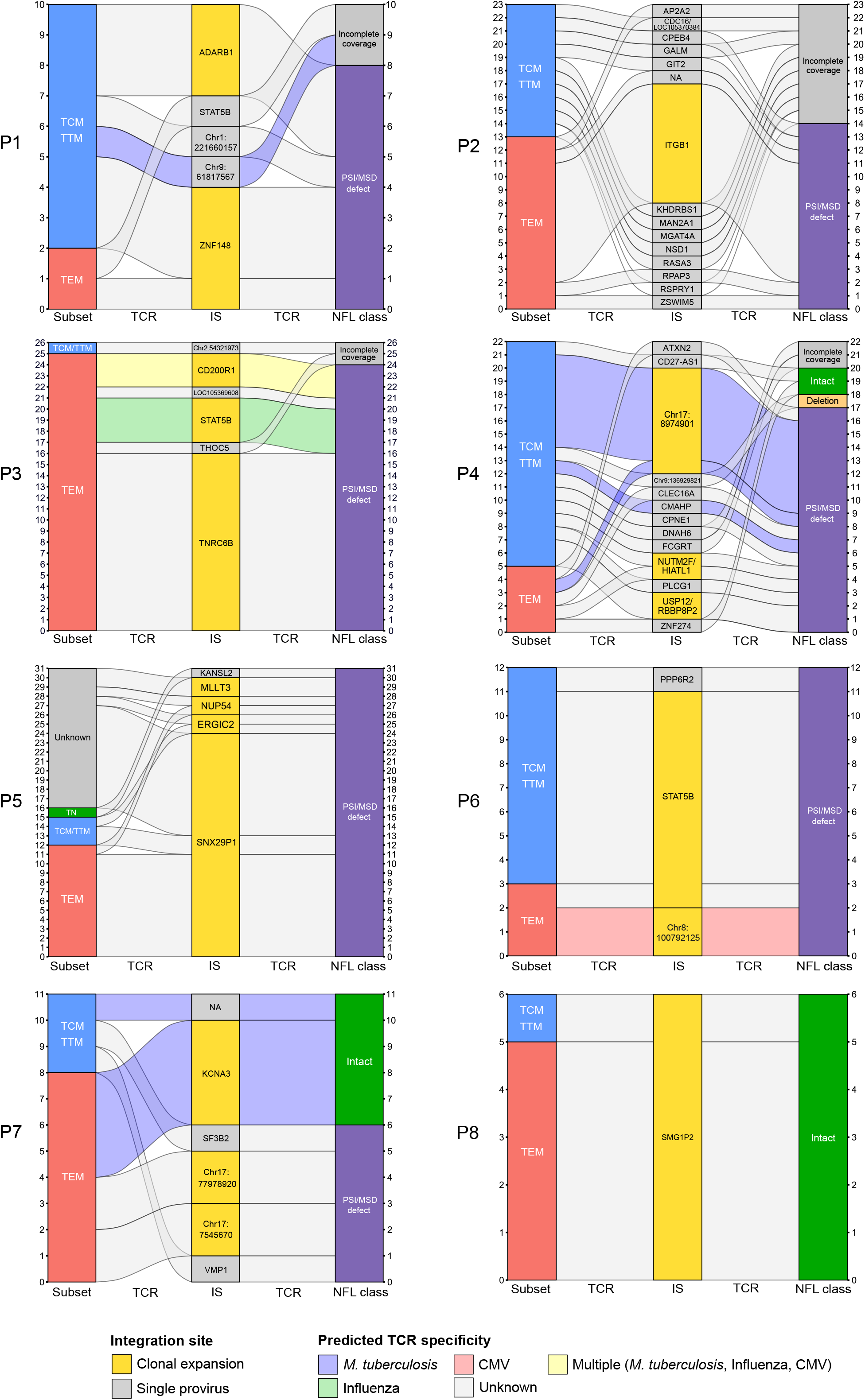
Predicted TCR specificity of single p24-producing cells. Alluvial plots showing the memory phenotype of the host cell, the IS and the NFL class for each p24-producing cell from n=8 ART-treated individuals. Single p24+ sorted cells are represented on the y-axis of each plot. Alluvials connecting the different categories are colored according to predicted TCR specificity. Only time points on ART are represented on this figure. IS = integration site, TCR = T cell receptor, NFL = near full-length, TN = naïve T cell, TCM = central memory T cell, TTM = transitional memory T cell, TEM = effector memory T cell.

Participant P3 had a clone with a predicted cross-reactive TCR (CMV, influenza, *M. tuberculosis*), for which the provirus was integrated in *CD200R1*, a gene not known to be involved in cell proliferation (Fig. 3). Participants P4 and P6 displayed clones with predicted specificity towards *M. tuberculosis* and CMV respectively (Fig. 3). Both clones harbored a provirus integrated at an intergenic region (chr17:8974901 and chr8:100792125, respectively), suggesting that their expansion was not driven by promoter insertion (Fig. 3). In contrast, we found several antigen-responsive cells with IS in genes involved in cell proliferation, as previously described by Simonetti *et al*.^18^. Participant P3 harbored a clone with an IS in *STAT5B,* potentially allowing for IS-driven proliferation. Moreover, the TCR specificity towards influenza suggests that the seasonal flu or vaccination might have contributed to the expansion of this clone (Fig. 3). Similarly, one expanded clone from participant P7 had predicted specificity towards *M. tuberculosis* and had an intact provirus integrated in *KCNA3*, a gene involved in T cell activation and proliferation^51^. Of note, this provirus was integrated in the same orientation as the gene, which could lead to aberrant transcription and subsequent disregulation of *KCNA3* expression.

Finally, to investigate the dynamics of the translation-competent reservoir, we performed STIP-Seq on two longitudinal samples from P5, collected 3 years apart (Supplementary Fig. 6, Supplementary Table 1). While the largest clone at the first timepoint (IS in *SNX29P1/P2*) was not retrieved 3 years later, one new clone emerged (IS in *LOC105369901*) and two clones persisted (IS in *ERGIC2* and *MLLT3*). These observations confirm that HIV-infected cell clones can persist, contract or expand over time^17,19,52^.

Taken together, we show that antigen-responsive cells can harbor inducible proviruses that are integrated in genes associated with cell proliferation, suggesting that antigen exposure and IS-driven mechanisms can synergize to favor the persistence of translation-competent reservoirs.

### Proviral sequences recovered with STIP-Seq match plasma virus sequences obtained during ART and upon ATI

We then investigated whether proviruses retrieved with STIP-Seq overlap with plasma virus sequences before and during an ATI. We performed STIP-Seq on CD4 T cells from three participants (P6, P7, P8), both during ART (T1; last time point before ATI) and during the ATI (T2; last available time point with undetectable viremia during ATI) (Fig. 4a, b). Plasma viral sequences (V1-V3 *env,* 894bp) from before (T1) and during the ATI (T2, T3, T4) were aligned to trimmed NFL sequences obtained with STIP-Seq, and maximum-likelihood phylogenetic trees were constructed (Fig. 5). The viral reservoir of two of the three participants (P6, P7) was previously characterized at T1 by FLIPS and *Matched Integration site and Proviral Sequencing* (MIP-Seq), providing an extensive resource for comparison with the STIP-Seq assay^26,53^. To this end, NFL proviral genomes obtained with FLIPS and MIP-Seq were also trimmed to the V1-V3 *env* region and included in the phylogenetic trees (Fig. 5).

**Figure 4:**
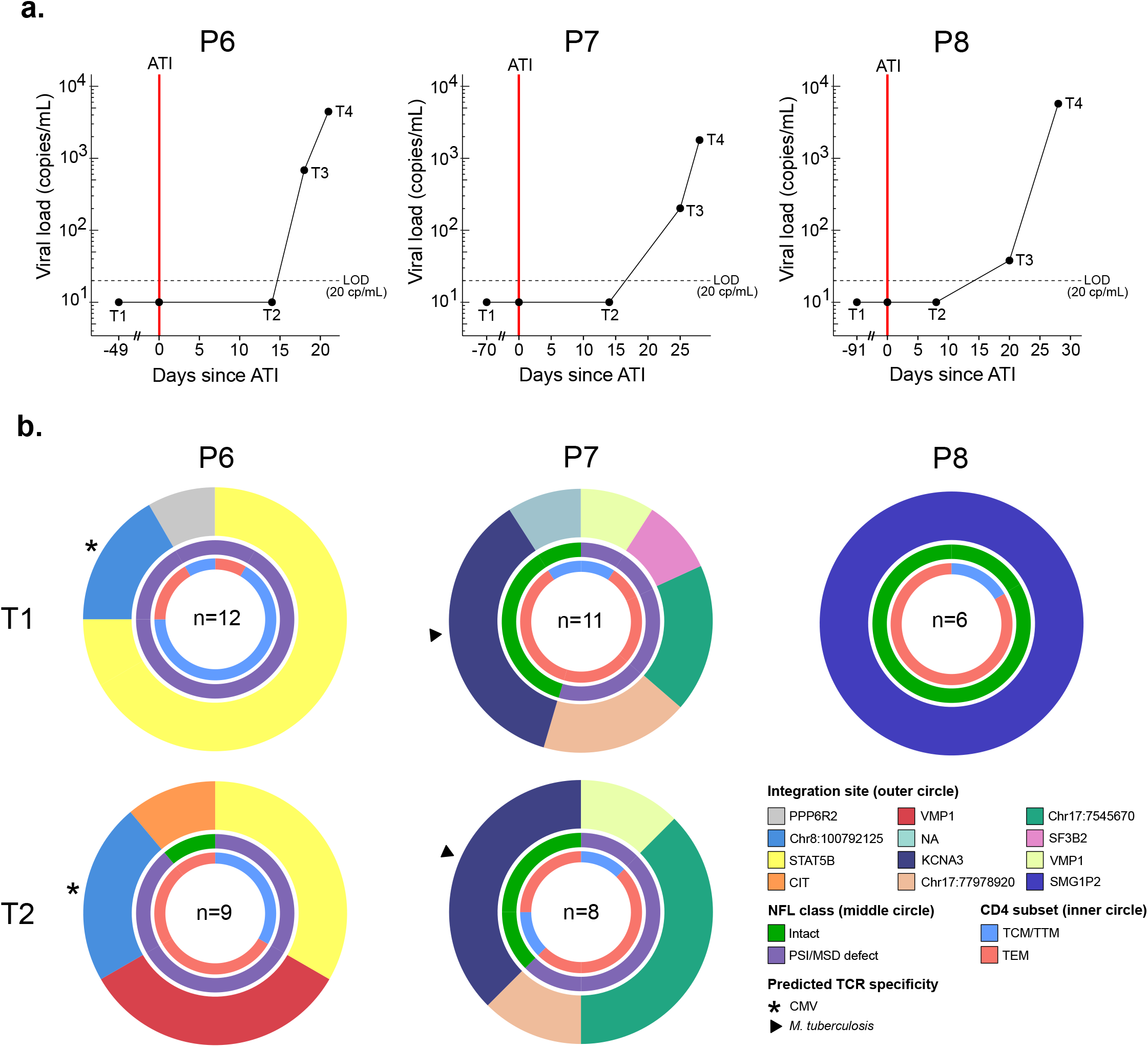
STIP-Seq on three participants before and during an analytical treatment interruption (ATI). (**a**) Viral load diagram showing the sampling timepoints before (T1) and during (T2, T3, T4) the analytical treatment interruption for 3 participants (P6, P7, P8). The viral load was undetectable at T1 and T2, under 1000 cp/mL at T3 (early rebound) and above 1000 cp/mL at T4 (late rebound). The vertical red line depicts the start of the ATI. LOD = limit of detection. (**b**) Donut charts displaying integration sites, NFL class, and memory subsets of p24-producing cells recovered before (T1) and during (T2) an ATI. The number of analyzed p24+ cells is indicated for each participant. PSI = packaging signal, MSD = major splice donor, TCM = central memory T cell, TTM = transitional memory T cell, TEM = effector memory T cell, NFL = near full-length.

**Figure 5:**
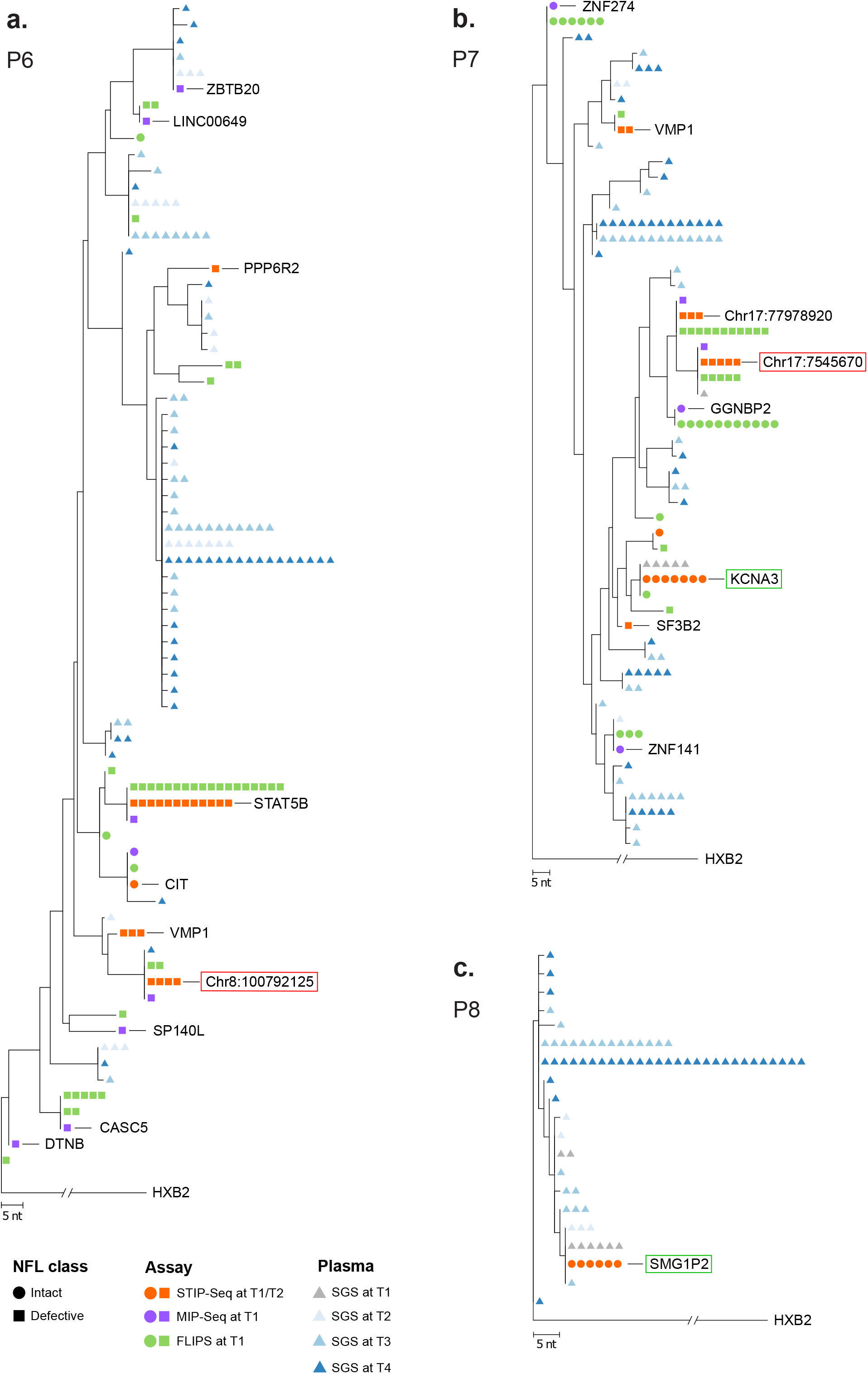
STIP-Seq identifies clones responsible for viremia under ART and upon treatment interruption. (**a-c**) Maximum-likelihood phylogenetic trees for 3 participants who underwent an analytical treatment interruption. The trees include V1-V3 *env* plasma sequences from before (T1) and during (T2, T3, T4) the treatment interruption (P6, P7, P8), as well as STIP-Seq, MIP-Seq and FLIPS sequences (T1) that were trimmed to the V1-V3 *env* region (P6, P7). Intact and defective proviruses are represented by circles and squares respectively, while V1-V3 plasma sequences are represented by triangles. Each assay is color-coded. Clones displaying a match between defective and intact STIP-Seq sequences and plasma sequences are indicated by red and green frames, respectively. HXB2 = subtype B HIV-1 reference genome, NFL = near full-length, STIP-Seq = Simultaneous TCR, Integration site and Provirus sequencing, MIP-Seq = Matched Integration site and Provirus sequencing, FLIPS = Full-Length Individual Provirus sequencing.

A total number of 29 p24+ cells at T1 and 17 p24+ cells at T2 were recovered (Fig. 4b). Overall, little differences were observed between the two timepoints, with most of the clones identified under ART (T1) persisting during the ATI (T2) (Fig. 4b). However, participant P6 displayed a novel clone at T2 when compared to T1, with an IS in the *VMP1* gene (Fig. 4b). Interestingly, the provirus from this clone did not match any V1-V3 *env* SGS, FLIPS, MIP-Seq or STIP-Seq sequences obtained at T1 (together evaluating n=382 proviruses) (Fig. 5a). In contrast, 3 out of 9 cells recovered by STIP-Seq at T2 yielded this provirus, indicating that this clone emerged or enlarged during the ATI.

For participant P6, one plasma sequence obtained during the ATI (T4) matched a provirus (IS at chr8:10079212) that was recovered with STIP-Seq at T1 (n=2) and T2 (n=2) (Fig. 5a, indicated with a red box). Interestingly, this provirus had a deletion at the 5’-end of the genome covering a large portion of the *p17* gene (Supplementary Fig. 3), making it unlikely that it could produce infectious virions. We previously established that the clonal prediction score (CPS) for the V1-V3 *env* region of participant P6 is 95% (based on n=22 NFL genomes with detectable V1-V3), indicating that while this score is high, this subgenomic region is not capable of differentiating all distinct proviruses^53^. Therefore, we cannot exclude the possibility that this plasma sequence stems from another provirus that has the same V1-V3 *env* sequence, though differs elsewhere in the genome.

For participant P7, five identical plasma sequences recovered at T1 matched an intact provirus (IS in *KCNA3*) that was identified with STIP-Seq at T1 (n=4) and T2 (n=3), indicating that this clone was responsible for low-level viremia (LLV) production under ART (Fig. 5b, indicated with a green box). Interestingly, this clone had a predicted TCR specificity against *M. tuberculosis,* suggesting that clones responsible for LLV on ART can proliferate in response to a circulating antigen (Fig. 4b). Similarly, one plasma sequence recovered during T1 matched a provirus (IS at chr17:7545670) that was identified with STIP-Seq at T1 (n=2) and T2 (n=3) (Fig. 4b). This provirus displayed a 5bp deletion in stem-loop 2 which removed the MSD, suggesting that a deletion of the MSD would still allow for detectable virion production (Fig. 5b, indicated by a red box). As calculated previously, the CPS for the V1-V3 *env* region of participant P7 is 100% (based on n=17 NFL genomes with detectable V1-V3), giving confidence about the validity of these matches^53^. Interestingly, FLIPS and MIP-Seq identified 3 additional proviruses that are genome-intact, but were not detected with STIP-Seq (IS in *ZNF274, ZNF141, GGNBP2*) (Fig. 5b). The low number of sampled p24+ cells (n=11 at T1, n=8 at T2, Fig. 4b) could potentially explain this observation, although is it also possible that these proviruses were not induced after a single round of PMA/ionomycin stimulation.

For participant P8, a single clone was identified, with a genome-intact provirus integrated in the *SMG1P2* pseudogene (Fig. 4b). The proviral sequence matched plasma sequences at T1, T2 and T3 (n=6, 3, and 1, respectively), suggesting that this clone was responsible for LLV production under ART, and further contributed to rebound viremia upon ATI (Fig. 5c, indicated by a green box). Because FLIPS data for this participant was not available, the CPS could not be calculated. Alternatively, the nucleotide diversity at T1 was calculated based on proviral V1-V3 *env* sequences, revealing a low diversity (0.00318 vs. 0.01579 for P6 and 0.01805 for P7)^26^, which could potentially lead to inaccurate links.

In conclusion, we show that STIP-Seq captures clones that contribute to LLV and viral rebound, and that in some cases, clones contributing to viral rebound already produce LLV during ART. Furthermore, our data suggest that clones responsible for LLV during ART can proliferate in response to antigenic stimulation.

## Discussion

HIV cure is impeded by the existence of a persistent viral reservoir, capable of refueling viremia upon treatment interruption. Unraveling mechanisms of viral latency and reservoir maintenance through clonal proliferation are research priorities in the field. Previous studies have shown that reservoir persistence is the result of a complex interplay between proviral genome integrity^28,30^, IS^22,33,50^ and antigenic stimulation of infected cells^16–18^, among other factors. In this regard, several assays have been developed to investigate these factors individually in ART-suppressed individuals. Here, we introduce a novel method to simultaneously characterize the NFL proviral genome and IS of translation-competent proviruses, as well as the phenotype and TCR sequence of the host cells. STIP-Seq requires only a limited amount of CD4 T cells (~5-10 million) and overcomes the need for limiting dilutions, as each sorted p24+ cell is HIV-infected. As a result, STIP-Seq is less labor and reagent intensive than MDA-based approaches on bulk DNA.

Conducting STIP-Seq on blood samples from 8 ART-suppressed individuals allowed for an in-depth characterization of the translation-competent reservoir. Only 12.5% of proviruses recovered with STIP-Seq were putatively intact, indicating that a large fraction of the translation-competent reservoir might not be replication-competent, as previously suggested^37,45^. Interestingly, a large proportion (45%) of the proviruses had intact open reading frames for all the protein coding genes, in contrast with proviruses obtained on bulk DNA, which often display large internal deletions, inversions or hypermutations^28,29^. Nevertheless, most of the proviruses recovered with STIP-Seq had small deletions (<500 bp) at the 5’-end of the genome, frequently involving a deletion of the MSD, as well as the cryptic donor (CD) site located 4 bp downstream of the MSD^30,54^. The presence of either of these sites was previously thought to be essential for correct splicing of viral transcripts and subsequent translation into viral proteins^54,55^. However, Pollack *et al.* showed that proviruses can bypass MSD deletions and mutations by activating alternative splice donor sites^56^. Here, we showed that proviruses with MSD/CD deletions can produce detectable amounts of p24 protein, suggesting that Tat/Rev mRNA can be produced despite MSD/CD mutations and/or that p24 production following PMA/ionomycin stimulation can happen in a Tat/Rev-independent manner. Indeed, an *in vitro* study showed that Tat-defective HIV-strains can produce readily detectable p24 following PMA stimulation^57^. Also, since it has been reported that PMA increases the expression of active NF-kB and P-TEFb^58^, and that NF-kB can directly bind P-TEFb to promote elongation of transcription in a Tat-independent manner^59^, it is likely that P-TEFb recruitment by NF-kB increases the levels of unspliced RNA.

We also found several translation-competent proviruses with deletions in the packaging signal (PSI), frequently spanning multiple stem-loops. Although this observation suggests that these proviruses are not replication-competent, previous studies have shown that viral genomes can still be packaged despite PSI defects, though with a considerably lower efficiency^56,60^. Importantly, it has been shown that MSD/PSI defective proviruses can produce viral proteins that can be recognized by cytotoxic CD8 T cells, leading to chronic immune activation^56,61^. We therefore conclude that while STIP-Seq does not solely enrich for genome-intact proviruses, it does enrich for proviruses that are potentially involved in HIV-1 pathogenesis. Future studies on MSD/PSI-defective proviruses will have to be conducted to further elucidate the effect of MSD/PSI deletions on replication-capacity, including a detailed assessment of viral splicing products and cloning of MSD/PSI-defective genomes into expression vectors.

We identified three distinct IS into *STAT5B*, a gene that was previously described as a hotspot for HIV-integration in ART-suppressed individuals^19–21,48^. A previous study has shown that integration in *STAT5B* in the same orientation as the gene can lead to aberrant splicing and subsequent cellular proliferation^21^. Interestingly, the 3 proviruses identified in the present study were integrated in the reverse orientation. However, studies that reported an overrepresentation of IS in the *STAT5B* gene showed that these IS could be found in both orientations, without an apparent bias^62^. This suggests that integration in the reverse orientation could still lead to clonal expansion, driven by other mechanisms than virus-host aberrant splicing.

Furthermore, we identified several infected cell clones with predicted specificities towards CMV, *M. tuberculosis* and influenza, underlining the role of antigen stimulation as a driver of clonal expansion^16–18^. In accordance with results from Simonetti *et al.*, we found antigen-responsive clones with IS in genes involved in cell proliferation (*STAT5B*, *KCNA3*), strengthening the hypothesis of a synergetic effect between IS-driven and antigen-driven proliferation^18^. Furthermore, our data suggest that one of these clones is responsible for LLV under ART (*KCNA3*), providing evidence that the proliferation of such clones can be driven by antigenic stimulation and/or IS-specific mechanisms.

In the context of an ATI, we compared p24+ cells obtained before (T1) and at the beginning of the ATI (T2). In one participant, a novel clone emerged during the ATI, which was not detected by V1-V3 *env* SGS, FLIPS, MIP-Seq, or STIP-Seq at T1. As we have previously shown that interferon-stimulated genes are already upregulated at T2 despite an undetectable viral load^63^, we hypothesize that this clonal expansion might have been driven by the inflammatory environment^64^. In addition, we found identical sequences between proviruses recovered with STIP-Seq and plasma viral sequences before and during the ATI, suggesting that HIV-infected clones can produce LLV during ART and/or contribute to rebound viremia upon ATI. This observation is in line with findings from Kearney *et al.*, which showed overlap between proviral p6-PR-RT sequences (DNA and cell-associated RNA, ~1540 bp) and plasma sequences obtained during TI^65^. Because the matching p6-PR-RT sequences were often clonal in nature, this prior study suggested that initial rebound could be fueled by clonally expanded populations of infected cells. Similarly, Aamer *et al.* identified links between plasma sequences recovered during TI and clonal C2-V5 *env* plasma sequences (~600 bp) that persisted for several years under treatment, supporting the notion that clonal cell populations that produce LLV under ART can contribute to viral rebound^66^. Using MDA-based NFL and ISA on bulk CD4 T cell DNA, Halvas *et al.* identified clonal populations of proviruses that could be linked to plasma sequences in non-suppressed individuals on ART, though their contribution to rebound viremia was not investigated^67^. In the present study, we provide deeper insights by identifying the phenotype and predicted TCR specificity of such clones and linking them to rebounding plasma sequences.

We acknowledge several limitations to this study. First, due to limited sample availability, we were not able to perform the viral outgrowth assay (VOA). Therefore, we could not evaluate the replication-competence of proviral sequences obtained with STIP-Seq by comparing them to sequences from positive VOA wells. Such comparison would have been particularly interesting for the participants that underwent an ATI, given the notoriously poor overlap between sequences derived from VOA and rebound plasma sequences^68–72^. Next, the link to rebound plasma sequences was based on a subgenomic region of the viral genome (V1-V3 *env*). It has previously been shown that using a subgenomic region to link viral sequences is not always adequate, as some viruses share the same subgenomic sequence while differing elsewhere in the genome^34,73^. However, we previously determined the CPS for two of the three participants that underwent an ATI, revealing high scores: 95% for P6 and 100% for P7^53^. Although the CPS should not be considered definitive, these scores give confidence about the validity of the observed matches. Finally, like other assays based on reactivation of proviruses with a latency reversal agent (LRA), STIP-Seq probably does not pick up all translation-competent proviruses, as reactivation is a stochastic process^36,58^. In this regard, it has been suggested that the IS can have an influence on the reactivation of the provirus^33,50,74^, and that different LRAs might induce reactivation of distinct proviral species^75,76^. As such, future studies with STIP-Seq investigating the relationship between different classes of LRA and the IS of the reactivated proviruses, would be of great interest.

In conclusion, our STIP-Seq assay enables deep characterization of the translation-competent HIV reservoir by simultaneously capturing four layers of information: NFL proviral genome, IS, phenotype and the TCRβ sequence of the host cell. By conducting this assay on ART-suppressed individuals, we provide further insights on the composition of the translation-competent reservoir and its persistence by clonal proliferation. Applying STIP-Seq in the context of an ATI revealed that cell clones harboring translation-competent proviruses contribute to residual viremia and viral rebound upon ART interruption. Using STIP-Seq on a larger cohort of individuals, along with a more elaborate panel of antibodies and different types of LRAs, will help to further unravel the complex interplay between viral and cellular factors involved in the long-term persistence of the HIV reservoir.

## Methods

### Participants and blood collection

A total of 8 individuals on stably suppressive ART were included in this study (Supplementary Table 1). Participants P1-P4 were recruited at the McGill University Health Centre and the Centre Hospitalier de l’Université de Montréal. Participants P5-P8 were recruited at Ghent University Hospital. Participants P6-P8 are part of the HIV-STAR cohort (Ghent University) (P6 = STAR 10, P7 = STAR 11, P8 = STAR 3). All participants underwent leukapheresis to collect large numbers of PBMCs. PBMCs were isolated by Ficoll density gradient centrifugation and were cryopreserved in liquid nitrogen.

### Ethics statement

All participants were adults and signed informed consent forms approved by the Ethics Committee of the Ghent University Hospital (Belgium), McGill University Health Centre and Centre Hospitalier de l’Université de Montréal (Canada).

### Antibodies

Fixable Viability Stain 510 was obtained from ThermoFisher Scientific (L34957). The following antibodies were used in sorting experiments: CD8 AF700 Clone RPA-T8 (ThermoFisher, 56-0088-41), CD45RO BV421 Clone UCHL1 (BD Biosciences, 562649), CD27 BV605 Clone L128 (BD Biosciences, 562656). For p24 staining, we used a combination of two antibodies: p24 KC57-FITC (Beckman Coulter, 6604665) and p24 28B7-APC (MediMabs, MM-0289-APC).

### Negative selection of CD4 T cells

CD4 T cells were isolated from PBMC by negative magnetic selection using the EasySep Human CD4 T Cell Enrichment Kit (StemCell Technology, 19052). Purity was typically >98%.

### HIV-Flow procedure

5-10×10^6^ CD4 T cells were resuspended at 2×10^6^ cells/mL in RPMI + 10% Fetal Bovine Serum and antiretroviral drugs were added to the culture (200nM raltegravir, 200nM lamivudine) to avoid new cycles of replication. Cells were stimulated with 1μg/mL ionomycin (Sigma, I9657) and 162nM PMA for 24h (Sigma, P8139). Frequencies of p24+ cells were measured by using a combination of 2 antibodies targeting the p24 protein (p24 KC57-FITC, p24 28B7-APC) as previously described by Pardons *et al*.^41^

### Methanol-based HIV-Flow procedure (STIP-Seq)

5-10×10^6^ CD4 T cells were resuspended at 2×10^6^ cells/mL in RPMI + 10% Fetal Bovine Serum (FBS, HyClone RB35947) and antiretroviral drugs were added to the culture (200nM raltegravir, 200nM lamivudine) to avoid new cycles of replication. Cells were stimulated with 1μg/mL ionomycin (Sigma, I9657) and 162nM PMA (Sigma, P8139). After a 24h-stimulation, a maximum of 10×10^6^ cells per condition were resuspended in PBS and stained with fixable viability stain 510 for 20 min at RT. Cells were then stained with antibodies against cell surface molecules (CD8, CD45RO, CD27) in PBS + 2% FBS for 20min at 4°C. After a 5 min-centrifugation step at 4°C to pre-chill the cells, CD4 cells were vortexed to avoid clumping and 1mL of ice-cold methanol (−20°C) was gently added. Cells were fixed/permeabilized in methanol for 15 min on ice. Intracellular p24 staining was performed in PBS + 2% FBS using a combination of 2 antibodies (p24 KC57-FITC, p24 28B7-APC) (45min, RT). Cells were then washed and resuspended in PBS for subsequent sorting. In all experiments, CD4 T cells from an HIV-negative control were included to set the threshold of positivity. The detailed protocol of the methanol-based HIV-Flow procedure can be found here: https://protocols.io/view/methanol-based-hiv-flow-bpedmja6.

### Single cell sorting of p24+ cells by fluorescence-activated cell sorting (FACS)

Single p24+ cells were sorted on a BD FACSAria™ Fusion Cell Sorter. The gating strategy used to sort the cells is represented in Supplementary Fig. 7. Cells were sorted in skirted 96-well PCR plates (Biorad, Cat. No. 12001925), into a volume of 4 μL PBS sc 1X (Qiagen, Cat. No. 150345). To avoid evaporation of the PBS sc 1X during the sort, the PCR plate was continuously chilled at 4°C. Index sorting was used to enable phenotyping of single sorted cells. CD4 T cell memory subsets were defined as follows: TN = CD45RO-CD27+, TCM/TTM = CD45RO+ CD27+, TEM = CD45RO+ CD27−, TTD = CD45RO-CD27− (Supplementary Fig. 7). Flow-Jo software v10.6.2 was used to analyze flow cytometry data (Tree-Star).

### Multiple Displacement Amplification (MDA)

Whole genome amplification of single sorted cells was carried out by multiple displacement amplification with the REPLI-g single cell kit (Qiagen, Cat. No. 150345), according to manufacturer’s instructions. A positive control, consisting of ten p24-cells sorted into the same well, was included on every plate.

### Quantitative polymerase chain reaction (qPCR) for RPP30

After whole genome amplification, reactions were screened by a binary qPCR on the RPP30 reference gene. The PCR mix consisted of 5 μL 2X LightCycler® 480 Probes Master (Roche, Cat. No. 04707494001), 1 μL MDA product, 0.4 μL 10 μM forward primer (5’-AGATTTGGACCTGCGAGCG-3’), 0.4 μL 10 μM reverse primer (5’-GAGCGGCTGTCTCCACAAGT-3’), 0.2 μL 10 μM probe (5’-TTCTGACCTGAAGGCTCTGCGCG-3’) and 3 μL nuclease free water. Reactions that yielded a cycle of threshold (Ct) value of 38 or lower, were selected for further downstream processing (Supplementary Figure 2).

### Integration site analysis

MDA reactions that were positive for RPP30 were subjected to integration site sequencing by a modified version of the integration site loop amplification (ISLA) assay, as described^53^. Resulting amplicons were visualized on a 1% agarose gel and positives were sequenced by Sanger sequencing. Analysis of the sequences was performed using the ‘Integration Sites’ webtool (https://indra.mullins.microbiol.washington.edu/integrationsites). Cancer-related genes were identified as described previously^77,78^.

### Near Full-Length proviral sequencing

Near full-length HIV-1 proviral sequencing was performed on MDA wells that were RPP30 positive (Supplementary Fig. 1). First, a set of five non-multiplexed PCRs was used to amplify the proviral genome, yielding five amplicons of approximately 2 kb in length that together cover 92% of the HIV-1 genome, as described^33^. Amplicons were visualized on a 1% agarose gel. MDA wells that did not yield an amplicon for all five PCRs were subjected to left and right half genome amplification. The 25 μL PCR mix for the first round is composed of: 5 μL 5X Prime STAR GXL buffer, 0.5 μL PrimeStar GXL polymerase (Takara Bio, Cat. No. R050B), 0.125 μL ThermaStop (Sigma Aldrich, Cat. No. TSTOP-500) 250 nM forward primer, 250 nM reverse primer and 1 μL REPLI-g product. The mix for the second round has the same composition and takes 1 μL of the first-round product as an input. Thermocycling conditions for first and second PCR rounds are as follows: 2 min at 98°C; 35 cycles (10 sec at 98°C, 15 sec at 62°C, 5 min at 68°C); 7 min at 68°C. For selected wells, NFL amplification using a set of four non-multiplexed PCRs was performed, as described^34^. The primer sequences for the five-, two- and four-amplicon approaches are summarized in Supplementary Table 5. Amplicons were pooled and cleaned by magnetic bead purification (Ampure XP, Beckman Coulter, Cat. No. A63881). Library preparation and sequencing was performed by short-read Illumina sequencing, as described^28^. De-novo HIV-1 genome assembly was performed as described^53^. Intactness classification was performed manually, using the criteria described by Pinzone *et al.*^30^

### Full-Length Individual Provirus Sequencing (FLIPS) on bulk CD4 T cell DNA

Full-Length Individual Provirus Sequencing was performed on DNA extracted from total CD4 T cells with the DNeasy Blood & Tissue Kit (Qiagen, Cat. No. 69504), as described^28^. Intactness classification was performed manually, using the criteria described by Pinzone *et al.*^30^

### Phylogenetic analyses

Sequences obtained with STIP-Seq, MIP-Seq and FLIPS were trimmed to the V1-V3 *env* region and multiple aligned to V1-V3 *env* sequences from plasma using MAFFT^79^. Phylogenetic trees were constructed using PhyML v3.0 (best of NNI and SPR rearrangements) and 1000 bootstraps^80^. MEGA7 and iTOL v5 were used to visualize phylogenetic trees^81,82^.

### TCR sequencing

A previously developed two-step PCR method to amplify a portion of approximately 260bp of the TCRβ encompassing (including the CDR3 region) was applied to MDA positive wells^17^. Briefly, a first multiplex PCR was performed using a set of 35 primers. M13 forward and reverse tags were included to the 5’ end of these primers, to allow a second PCR amplification, which was followed by Sanger sequencing, with M13F and M13R as sequencing primers. TCRβ sequences were re-constructed using both forward and reverse sequences, and were analyzed using the V-QUEST tool of the IMGT® database (IMGT®, the international ImMunoGeneTics information system® [http://www.imgt.org])^83^.

### Prediction of TCR specificity

TCR sequences were analyzed using an algorithm to predict antigen specificity: CDR3 sequences were compared to the McPAS-TCR database of TCRs of known antigenic specificity ([http://friedmanlab.weizmann.ac.il/McPAS-TCR/], PMID: 28481982) and sequence similarities were identified. We predicted TCR specificity using the three criteria described by Meysman *et al.*^84^: 1) CDR3 sequences should have identical length, 2) CDR3 sequences should be long enough and 3) CDR3 sequences should not differ by more than one amino acid. Among all CDR3 sequences, those fulfilling these three criteria with matched CDR3 sequences from the database were considered at high probability of sharing the same specificity.

### Data representations and statistical analyses

Bar charts, line plots and donut plots were generated in R (version 3.4.3) or Graphpad Prism (version 8.0.2). Alluvial plots were generated in R (version 3.4.3) using the ggalluvial package (version 0.12.3). For group comparisons, non-parametric Wilcoxon matched-pairs signed rank tests were used. P values of less or equal to 0.05 were considered statistically significant.

## Data availability

Data will be uploaded to public repositories upon acceptance of the manuscript.

## Acknowledgements and funding sources

We thank all participants who donated samples to the study, as well as MDs and study nurses who helped with the recruitment and coordination of this study and the processing of blood samples. The study team thanks Sophie Vermaut for assisting with the flow cytometry platform. We also thank Bram Parton, Céline Helsmoortel and Kim De Leeneer for helping with the Illumina sequencing. We are grateful for the interesting scientific input and technical help given by Rémi Fromentin, Caroline Dufour, Amélie Pagliuzza and Sofie Rutsaert. In addition, we thank Jean-Pierre Routy and Josée Girouard for the recruitment of the participants in Montreal. This current research work was supported by the NIH (R01-AI134419, MPI: LV and JIM) and the Research Foundation Flanders (S000319N and G0B3820N). This work was partially supported by the Canadian Institutes for Health Research (CIHR; operating grant #364408 and the Canadian HIV Cure Enterprise (CanCURE) Team Grant HB2 - 164064). BC was supported by FWO Vlaanderen (1S28918N). LL was supported by FWO Vlaanderen (1S29220N). LV was supported by the Research Foundation Flanders (1.8.020.09.N.00) and the Collen-Francqui Research Professor Mandate. MP was supported by postdoctoral funding from VLAIO O&O (HBC.2018.2278). PG was supported by a postdoctoral fellowship from CIHR (#415209), and NC was supported by Research Scholar Career Awards of the FRQ-S (#253292). SP was supported by the Delaney AIDS Research Enterprise (DARE) to Find a Cure (1U19AI096109 and 1UM1AI126611-01) and the Australian National Health and Medical Research Council (APP1061681 and APP1149990).

## Author contributions

BC, MP, LL, WW, NC and LV conceptualized the experiments. Additional scientific input was given by NC, SP, WW and LV. NC and LV provided the samples used in the study. BC, MP, LL, YN and NB performed experiments involving cell sorting, multiple displacement amplification, single NFL proviral sequencing and integration site sequencing. SP provided protocols and resources for FLIPS sequencing. PG performed TCR sequencing. JIM and LC provided protocols to perform the 2-amplicon PCR for NFL proviral sequencing. BC and MP wrote the paper. All authors read and edited the paper.

## References

1. Finzi, D. et al. Identification of a reservoir for HIV-1 in patients on highly active antiretroviral therapy. Science 278, 1295–300 (1997).

2. Wong, J. K. et al. Recovery of replication-competent HIV despite prolonged suppression of plasma viremia. Science 278, 1291–1295 (1997).

3. Chun, T. W. et al. Presence of an inducible HIV-1 latent reservoir during highly active antiretroviral therapy. Proc. Natl. Acad. Sci. U. S. A. 94, 13193–13197 (1997).

4. Chun, T. W. et al. Early establishment of a pool of latently infected, resting CD4(+) T cells during primary HIV-1 infection. Proc. Natl. Acad. Sci. U. S. A. 95, 8869–73 (1998).

5. Finzi, D. et al. Latent infection of CD4+ T cells provides a mechanism for lifelong persistence of HIV-1, even in patients on effective combination therapy. Nat. Med. 5, 512–517 (1999).

6. Chun, T.-W. et al. Rebound of plasma viremia following cessation of antiretroviral therapy despite profoundly low levels of HIV reservoir: implications for eradication. AIDS Lond. Engl. 24, 2803–2808 (2010).

7. Colby, D. J. et al. Rapid HIV RNA rebound after antiretroviral treatment interruption in persons durably suppressed in Fiebig I acute HIV infection. Nat. Med. 24, 923–926 (2018).

8. Siliciano, J. D. et al. Long-term follow-up studies confirm the stability of the latent reservoir for HIV-1 in resting CD4+ T cells. Nat. Med. 9, 727–728 (2003).

9. Crooks, A. M. et al. Precise Quantitation of the Latent HIV-1 Reservoir: Implications for Eradication Strategies. J. Infect. Dis. 212, 1361–1365 (2015).

10. Chomont, N. et al. HIV reservoir size and persistence are driven by T cell survival and homeostatic proliferation. Nat. Med. 15, 893–900 (2009).

11. Bosque, A., Famiglietti, M., Weyrich, A. S., Goulston, C. & Planelles, V. Homeostatic proliferation fails to efficiently reactivate HIV-1 latently infected central memory CD4+ T cells. PLoS Pathog. 7, e1002288 (2011).

12. Vandergeeten, C. et al. Interleukin-7 promotes HIV persistence during antiretroviral therapy. Blood 121, 4321–4329 (2013).

13. Chomont, N., DaFonseca, S., Vandergeeten, C., Ancuta, P. & Sékaly, R.-P. Maintenance of CD4+ T-cell memory and HIV persistence: keeping memory, keeping HIV. Curr. Opin. HIV AIDS 6, 30–36 (2011).

14. Kumar, N. A. et al. Antibody-Mediated CD4 Depletion Induces Homeostatic CD4+ T Cell Proliferation without Detectable Virus Reactivation in Antiretroviral Therapy-Treated Simian Immunodeficiency Virus-Infected Macaques. J. Virol. 92, e01235–18 (2018).

15. Bacchus-Souffan, C. et al. Relationship between CD4 T cell turnover, cellular differentiation and HIV persistence during ART. PLoS Pathog. 17, e1009214 (2021).

16. Mendoza, P. et al. Antigen-responsive CD4+ T cell clones contribute to the HIV-1 latent reservoir. J. Exp. Med. 217, e20200051 (2020).

17. Gantner, P. et al. Single-cell TCR sequencing reveals phenotypically diverse clonally expanded cells harboring inducible HIV proviruses during ART. Nat. Commun. 11, 4089 (2020).

18. Simonetti, F. R. et al. Antigen-driven clonal selection shapes the persistence of HIV-1 infected CD4+ T cells in vivo. J. Clin. Invest. (2020) doi:10.1172/JCI145254.

19. Wagner, T. A. et al. HIV latency. Proliferation of cells with HIV integrated into cancer genes contributes to persistent infection. Science 345, 570–573 (2014).

20. Maldarelli, F. et al. Specific HIV integration sites are linked to clonal expansion and persistence of infected cells. Science 345, 179–183 (2014).

21. Cesana, D. et al. HIV-1-mediated insertional activation of STAT5B and BACH2 trigger viral reservoir in T regulatory cells. Nat. Commun. 8, 498 (2017).

22. Liu, R. et al. Single-cell transcriptional landscapes reveal HIV-1–driven aberrant host gene transcription as a potential therapeutic target. Sci. Transl. Med. 12, eaaz0802 (2020).

23. Hughes, S. H. & Coffin, J. M. What Integration Sites Tell Us about HIV Persistence. Cell Host Microbe 19, 588–598 (2016).

24. Mullins, J. I. & Frenkel, L. M. Clonal Expansion of Human Immunodeficiency Virus-Infected Cells and Human Immunodeficiency Virus Persistence During Antiretroviral Therapy. J. Infect. Dis. 215, S119–S127 (2017).

25. Liu, R., Simonetti, F. R. & Ho, Y.-C. The forces driving clonal expansion of the HIV-1 latent reservoir. Virol. J. 17, 4 (2020).

26. De Scheerder, M.-A. et al. HIV Rebound Is Predominantly Fueled by Genetically Identical Viral Expansions from Diverse Reservoirs. Cell Host Microbe 26, 347–358 (2019).

27. Lambrechts, L., Cole, B., Rutsaert, S., Trypsteen, W. & Vandekerckhove, L. Emerging PCR-Based Techniques to Study HIV-1 Reservoir Persistence. Viruses 12, (2020).

28. Hiener, B. et al. Identification of Genetically Intact HIV-1 Proviruses in Specific CD4+ T Cells from Effectively Treated Participants. Cell Rep. 21, 813–822 (2017).

29. Lee, G. Q. et al. Clonal expansion of genome-intact HIV-1 in functionally polarized Th1 CD4+ T cells. J. Clin. Invest. 127, 2689–2696 (2017).

30. Pinzone, M. R. et al. Longitudinal HIV sequencing reveals reservoir expression leading to decay which is obscured by clonal expansion. Nat. Commun. 10, 728 (2019).

31. Rousseau, C. M. et al. Large-scale amplification, cloning and sequencing of near full-length HIV-1 subtype C genomes. J. Virol. Methods 136, 118–125 (2006).

32. Cohn, L. B. et al. HIV-1 Integration Landscape during Latent and Active Infection. Cell 160, 420–432 (2015).

33. Einkauf, K. B. et al. Intact HIV-1 proviruses accumulate at distinct chromosomal positions during prolonged antiretroviral therapy. J. Clin. Invest. 129, 988–998 (2019).

34. Patro, S. C. et al. Combined HIV-1 sequence and integration site analysis informs viral dynamics and allows reconstruction of replicating viral ancestors. Proc. Natl. Acad. Sci. U. S. A. 116, 25891–25899 (2019).

35. Bruner, K. M. et al. Defective proviruses rapidly accumulate during acute HIV-1 infection. Nat. Med. 22, 1043–1049 (2016).

36. Ho, Y.-C. et al. Replication-competent noninduced proviruses in the latent reservoir increase barrier to HIV-1 cure. Cell 155, 540–551 (2013).

37. Abdel-Mohsen, M. et al. Recommendations for measuring HIV reservoir size in cure-directed clinical trials. Nat. Med. 26, 1339–1350 (2020).

38. Laird, G. M. et al. Rapid quantification of the latent reservoir for HIV-1 using a viral outgrowth assay. PLoS Pathog. 9, e1003398 (2013).

39. Lorenzi, J. C. C. et al. Paired quantitative and qualitative assessment of the replication-competent HIV-1 reservoir and comparison with integrated proviral DNA. Proc. Natl. Acad. Sci. U. S. A. 113, E7908–E7916 (2016).

40. Grau-Expósito, J. et al. A Novel Single-Cell FISH-Flow Assay Identifies Effector Memory CD4+ T cells as a Major Niche for HIV-1 Transcription in HIV-Infected Patients. mBio 8, e00876–17 (2017).

41. Pardons, M. et al. Single-cell characterization and quantification of translation-competent viral reservoirs in treated and untreated HIV infection. PLOS Pathog. 15, e1007619 (2019).

42. Baxter, A. E. et al. Single-Cell Characterization of Viral Translation-Competent Reservoirs in HIV-Infected Individuals. Cell Host Microbe 20, 368–380 (2016).

43. Cohn, L. B. et al. Clonal CD4+ T cells in the HIV-1 latent reservoir display a distinct gene profile upon reactivation. Nat. Med. 24, 604–609 (2018).

44. Neidleman, J. et al. Phenotypic analysis of the unstimulated in vivo HIV CD4 T cell reservoir. eLife 9, e60933 (2020).

45. Baxter, A. E., O’Doherty, U. & Kaufmann, D. E. Beyond the replication-competent HIV reservoir: transcription and translation-competent reservoirs. Retrovirology 15, 18 (2018).

46. Kearse, M. G. & Wilusz, J. E. Non-AUG translation: a new start for protein synthesis in eukaryotes. Genes Dev. 31, 1717–1731 (2017).

47. Yilmaz, A., Bolinger, C. & Boris-Lawrie, K. Retrovirus translation initiation: Issues and hypotheses derived from study of HIV-1. Curr. HIV Res. 4, 131–139 (2006).

48. Ikeda, T., Shibata, J., Yoshimura, K., Koito, A. & Matsushita, S. Recurrent HIV-1 integration at the BACH2 locus in resting CD4+ T cell populations during effective highly active antiretroviral therapy. J. Infect. Dis. 195, 716–725 (2007).

49. Hamann, M. V. et al. Transcriptional behavior of the HIV-1 promoter in context of the BACH2 prominent proviral integration gene. Virus Res. 293, 198260 (2020).

50. Jiang, C. et al. Distinct viral reservoirs in individuals with spontaneous control of HIV-1. Nature 585, 261–267 (2020).

51. Kang, J.-A. et al. Epigenetic regulation of Kcna3-encoding Kv1.3 potassium channel by cereblon contributes to regulation of CD4+ T-cell activation. Proc. Natl. Acad. Sci. U. S. A. 113, 8771–8776 (2016).

52. Wang, Z. et al. Expanded cellular clones carrying replication-competent HIV-1 persist, wax, and wane. Proc. Natl. Acad. Sci. 115, E2575–E2584 (2018).

53. Cole, B. et al. In-depth characterization of HIV-1 reservoirs reveals links to viral rebound during treatment interruption. Preprint at https://www.biorxiv.org/content/10.1101/2021.02.04.429690v1. (2021) doi:10.1101/2021.02.04.429690.

54. Purcell, D. F. & Martin, M. A. Alternative splicing of human immunodeficiency virus type 1 mRNA modulates viral protein expression, replication, and infectivity. J. Virol. 67, 6365–6378 (1993).

55. Das, A. T., Pasternak, A. O. & Berkhout, B. On the generation of the MSD-Ѱ class of defective HIV proviruses. Retrovirology 16, 19 (2019).

56. Pollack, R. A. et al. Defective HIV-1 Proviruses Are Expressed and Can Be Recognized by Cytotoxic T Lymphocytes, which Shape the Proviral Landscape. Cell Host Microbe 21, 494–506.e4 (2017).

57. Luznik, L., Kraus, G., Guatelli, J., Richman, D. & Wong-Staal, F. Tat-independent replication of human immunodeficiency viruses. J. Clin. Invest. 95, 328–332 (1995).

58. Pardons, M., Fromentin, R., Pagliuzza, A., Routy, J.-P. & Chomont, N. Latency-Reversing Agents Induce Differential Responses in Distinct Memory CD4 T Cell Subsets in Individuals on Antiretroviral Therapy. Cell Rep. 29, 2783–2795.e5 (2019).

59. Barboric, M., Nissen, R. M., Kanazawa, S., Jabrane-Ferrat, N. & Peterlin, B. M. NF-κB Binds P-TEFb to Stimulate Transcriptional Elongation by RNA Polymerase II. Mol. Cell 8, 327–337 (2001).

60. Lever, A., Gottlinger, H., Haseltine, W. & Sodroski, J. Identification of a sequence required for efficient packaging of human immunodeficiency virus type 1 RNA into virions. J. Virol. 63, 4085–4087 (1989).

61. Imamichi, H. et al. Defective HIV-1 proviruses produce viral proteins. Proc. Natl. Acad. Sci. 117, 3704–3710 (2020).

62. Anderson, E. M. & Maldarelli, F. The role of integration and clonal expansion in HIV infection: live long and prosper. Retrovirology 15, 71 (2018).

63. De Scheerder, M.-A. et al. Evaluating predictive markers for viral rebound and safety assessment in blood and lumbar fluid during HIV-1 treatment interruption. J. Antimicrob. Chemother. 75, 1311–1320 (2020).

64. Reed, J. M., Branigan, P. J. & Bamezai, A. Interferon gamma enhances clonal expansion and survival of CD4+ T cells. J. Interferon Cytokine Res. Off. J. Int. Soc. Interferon Cytokine Res. 28, 611–622 (2008).

65. Kearney, M. F. et al. Origin of Rebound Plasma HIV Includes Cells with Identical Proviruses That Are Transcriptionally Active before Stopping of Antiretroviral Therapy. J. Virol. 90, 1369–1376 (2016).

66. Aamer, H. A. et al. Cells producing residual viremia during antiretroviral treatment appear to contribute to rebound viremia following interruption of treatment. PLoS Pathog. 16, e1008791 (2020).

67. Halvas, E. K. et al. HIV-1 viremia not suppressible by antiretroviral therapy can originate from large T cell clones producing infectious virus. J. Clin. Invest. 130, 5847–5857 (2020).

68. Lu, C.-L. et al. Relationship between intact HIV-1 proviruses in circulating CD4+ T cells and rebound viruses emerging during treatment interruption. Proc. Natl. Acad. Sci. U. S. A. 115, E11341–E11348 (2018).

69. Vibholm, L. K. et al. Characterization of Intact Proviruses in Blood and Lymph Node from HIV-Infected Individuals Undergoing Analytical Treatment Interruption. J. Virol. 93, (2019).

70. Bertagnolli, L. N. et al. Autologous IgG antibodies block outgrowth of a substantial but variable fraction of viruses in the latent reservoir for HIV-1. Proc. Natl. Acad. Sci. U. S. A. 117, 32066–32077 (2020).

71. Cohen, Y. Z. et al. Relationship between latent and rebound viruses in a clinical trial of anti – HIV-1 antibody 3BNC117. J. Exp. Med. 215, 2311–2324 (2018).

72. Salantes, D. B. et al. HIV-1 latent reservoir size and diversity are stable following brief treatment interruption. J. Clin. Invest. 128, 3102–3115 (2018).

73. Laskey, S. B., Pohlmeyer, C. W., Bruner, K. M. & Siliciano, R. F. Evaluating Clonal Expansion of HIV-Infected Cells: Optimization of PCR Strategies to Predict Clonality. PLOS Pathog. 12, e1005689 (2016).

74. Vansant, G. et al. The chromatin landscape at the HIV-1 provirus integration site determines viral expression. Nucleic Acids Res. 48, 7801–7817 (2020).

75. Chen, H.-C., Martinez, J. P., Zorita, E., Meyerhans, A. & Filion, G. J. Position effects influence HIV latency reversal. Nat. Struct. Mol. Biol. 24, 47–54 (2017).

76. Darcis, G. et al. An In-Depth Comparison of Latency-Reversing Agent Combinations in Various In Vitro and Ex Vivo HIV-1 Latency Models Identified Bryostatin-1+JQ1 and Ingenol-B+JQ1 to Potently Reactivate Viral Gene Expression. PLoS Pathog. 11, e1005063 (2015).

77. Berry, C. C. et al. INSPIIRED: Quantification and Visualization Tools for Analyzing Integration Site Distributions. Mol. Ther. Methods Clin. Dev. 4, 17–26 (2017).

78. Sherman, E. et al. INSPIIRED: A Pipeline for Quantitative Analysis of Sites of New DNA Integration in Cellular Genomes. Mol. Ther. Methods Clin. Dev. 4, 39–49 (2017).

79. Katoh, K. & Standley, D. M. MAFFT multiple sequence alignment software version 7: improvements in performance and usability. Mol. Biol. Evol. 30, 772–780 (2013).

80. Guindon, S. et al. New algorithms and methods to estimate maximum-likelihood phylogenies: assessing the performance of PhyML 3.0. Syst. Biol. 59, 307–321 (2010).

81. Kumar, S., Stecher, G. & Tamura, K. MEGA7: Molecular Evolutionary Genetics Analysis Version 7.0 for Bigger Datasets. Mol. Biol. Evol. 33, 1870–1874 (2016).

82. Letunic, I. & Bork, P. Interactive Tree Of Life (iTOL) v4: recent updates and new developments. Nucleic Acids Res. 47, W256–W259 (2019).

83. Lefranc, M. P. et al. IMGT, the international ImMunoGeneTics database. Nucleic Acids Res. 27, 209–212 (1999).

84. Meysman, P. et al. On the viability of unsupervised T-cell receptor sequence clustering for epitope preference. Bioinformatics 35, 1461–1468 (2019).

