## Supplemental Materials for "In-depth single-cell analysis of translation-competent HIV-1 reservoirs identifies cellular sources of plasma viremia"

### Supplementary Materials

**Supplementary Fig. 1: Frequency of p24+ cells after methanol fixation: linearity and comparison with PFA.** (a-b) Linearity of the p24+ frequency with STIP-seq. J1.1 cells were serially diluted in Jurkat cells, fixed/permeabilized with methanol and stained with HIV p24-specific KC57 and 28B7 antibodies. (a) FACS dots plots of co-staining KC57/28B7 cells (b) Linear regression of p24+ frequency from serially diluted J1.1 cells in Jurkat cells. Predicted curve is represented by the dashed line. (c) Comparison of p24+ frequency using PFA-based fixation vs. methanol-based fixation on CD4 T cells from 1 viremic and 4 ART-treated individuals (P5 T1-T2, P6 T1, P7 T1, P8 T1). Each participant is represented by a color-coded symbol. For statistical analysis, a non-parametric matched-pairs Wilcoxon signed-rank test was used to compare frequencies. PFA = paraformaldehyde.

**Supplementary Fig. 2: STIP-Seq decision tree.** Step-by-step overview of the methodologies applied during STIP-Seq. Single sorted p24+ cells are subjected to whole genome amplification by multiple displacement amplification. Reactions are screened for successful amplification with a duplex qPCR (RPP30 reference gene, HIV LTR). Wells containing a cell with successfully amplified DNA are subjected to integration site analysis by integration site loop amplification, near full-length proviral sequencing with a 5- or 2-amplicon approach and TCR sequencing. WGA = whole genome amplification, LTR = long terminal repeat, ISLA = Integration Site Loop Amplification, NFL = Near full-length sequencing, TCR = T cell receptor.

**Supplementary Fig. 3: Deletions in the 5'UTR, p17 gene and p24 gene of translation-competent proviruses.** Graphical representation of the 5'UTR-p17-p24 region of proviruses recovered with STIP-Seq that had deletions spanning into p17/p24. Participant IDs are indicated on the left and the corresponding integration sites on the right side of each genome depiction. UTR = untranslated region.

**Supplementary Fig. 4: Comparison between FLIPS and STIP-Seq on 2 longitudinal samples from an ART-suppressed individual.** Maximum-likelihood phylogenetic trees from near full-length proviral genomes generated with FLIPS and STIP-Seq. STIP-Seq and FLIPS were performed on CD4 T cells from an ART-suppressed individual (P5), at two time points, 3 years apart. Bars depict the length of each proviral sequence, colored according to the NFL class. The blue arc depicts identical proviral genome sequences that were found with both FLIPS and STIP-Seq.

**Supplementary Fig. 5: Influence of methanol-fixation and PMA/ionomycin stimulation on memory subsets.** Representative FACS dot plots showing the co-staining of cells with CD45RO and CD27 (gated on live CD4 T cells). Purified CD4 T cells were stimulated or not with PMA/ionomycin and were subjected to either paraformaldehyde or methanol fixation after 24h. The central/transitional memory T cell quadrant (CD45RO+ CD27+; TCM/TTM) is colored in blue, the effector memory T cell quadrant (CD45RO+ CD27-; TEM) in red, the naïve T cell quadrant (CD45RO- CD27+; TN) in green and the terminally differentiated memory T cell quadrant (CD45RO- CD27-; TTd) in purple. PMA = phorbol myristate acetate.

**Supplementary Fig. 6: Longitudinal STIP-Seq analysis on ART-suppressed individual.** Alluvial plots showing the memory phenotype of the host cell, the IS and the NFL class for each p24-producing cell from two longitudinal samples of participant P5 (3 years apart). Single p24+ sorted cells are represented on the y-axis of each plot. IS = integration site, NFL = near full-length class, TN = naïve T cell, TCM = central memory T cell, TTM = transitional memory T cell, TEM = effector memory T cell.

**Supplementary Fig. 7: Gating strategy used in STIP-seq.** Representative example of the gating strategy used in STIP-Seq following PMA/ionomycin stimulation of CD4 T cells obtained from an ART-suppressed individual. p24+ (KC57+/28B7+) were single cell sorted. Index cell sorting allowed for *post hoc* analysis of the memory phenotype of p24+ cells, by using the CD27 and CD45RO markers. Arrows indicate the sequential gating order. FSC = forward scatter, SSC = side scatter, TN = naïve CD4 T cell, TCM/TTM = central/transitional memory CD4 T cell, TEM = effector memory CD4 T cell, TTD = terminally differentiated CD4 T cell.

**Supplementary Table 1: Clinical and virological characteristics of participants.** Participant P5 was sampled at two timepoints during ART-suppression (3 years apart), indicated in the Timepoint column as 'First' and '+3 years' respectively. Participants P6 and P7 were sampled during ART-suppression and during analytical treatment interruption, indicated in the Timepoint column as 'T1' and 'T2' respectively. ART = antiretroviral therapy; VL = viral load.

**Supplementary Table 2: Near-full length proviral PCRs.** Summary of results using the 5-amplicon, 2-amplicon and 4-amplicon PCR approaches for near full-length (NFL) proviral genome sequencing. The names of the amplicons are indicated above each column, with the HXB2 genome coordinates between parentheses. Green color = positive PCR result, red color = negative PCR result, grey color = not attempted. Frag = fragment, NA = not available, HXB2 = subtype B HIV-1 reference genome.

**Supplementary Table 3: Integration sites of p24+ cells.** Integration sites were mapped to the GRCh38.p2 human genome reference assembly. Cancer-related genes are indicated with an asterisk (*).  IS only retrieved at T2, during analytical treatment interruption, are indicated with a dagger (†). Chrom. = chromosome, NA = not applicable.

**Supplementary Table 4: TCR sequences and predicted specificities of p24+ cells.** TRBV/TRBJ usage and CDR3 amino acid sequence for each sorted p24+ cell. NA = not available.

**Supplementary Table 5: Primers used for near full-length proviral sequencing.** NFL = near full-length proviral sequencing, HXB2 = subtype B HIV-1 reference genome.

#### Supplementary Fig. 1

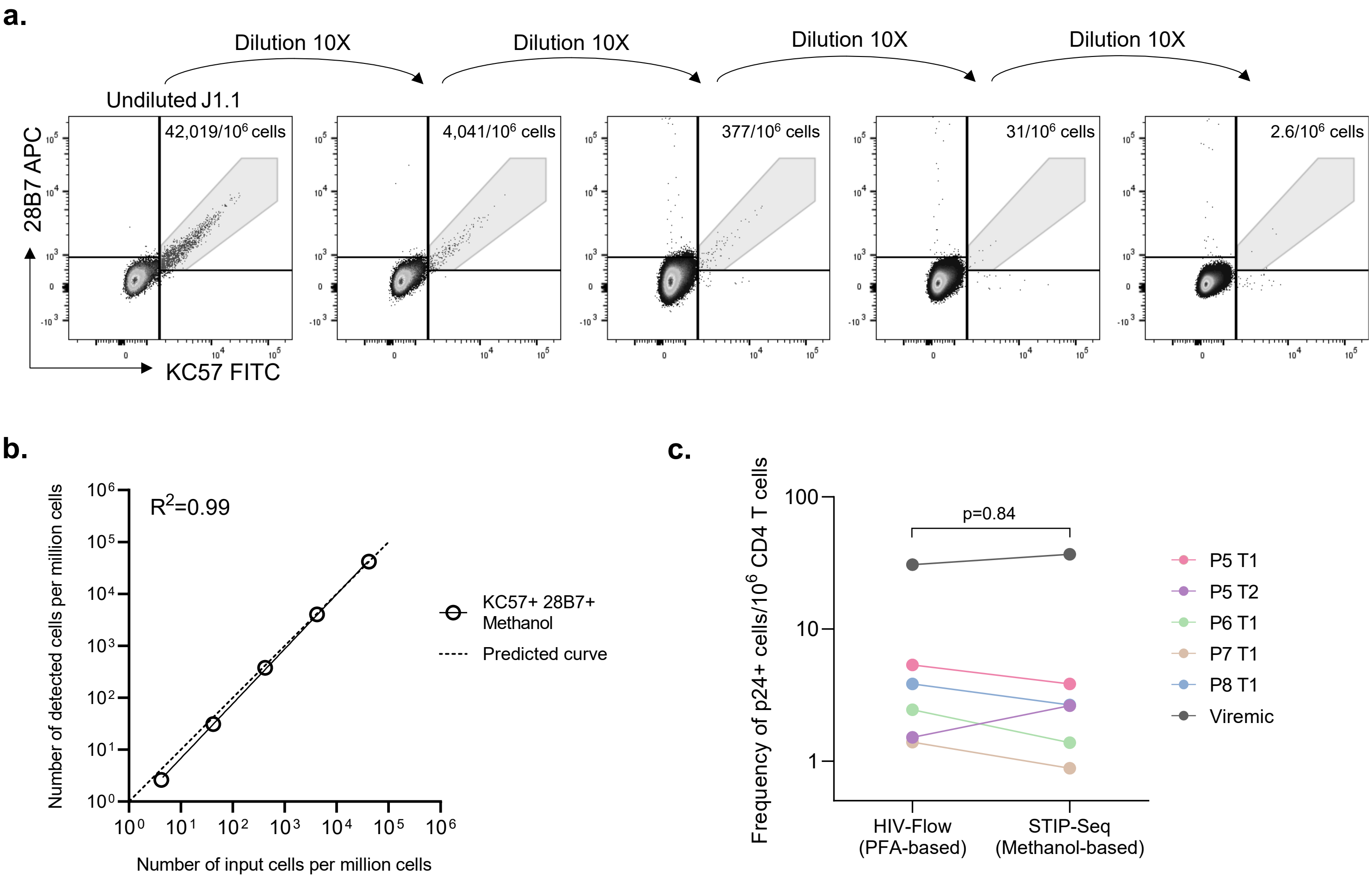

#### Supplementary Fig. 2

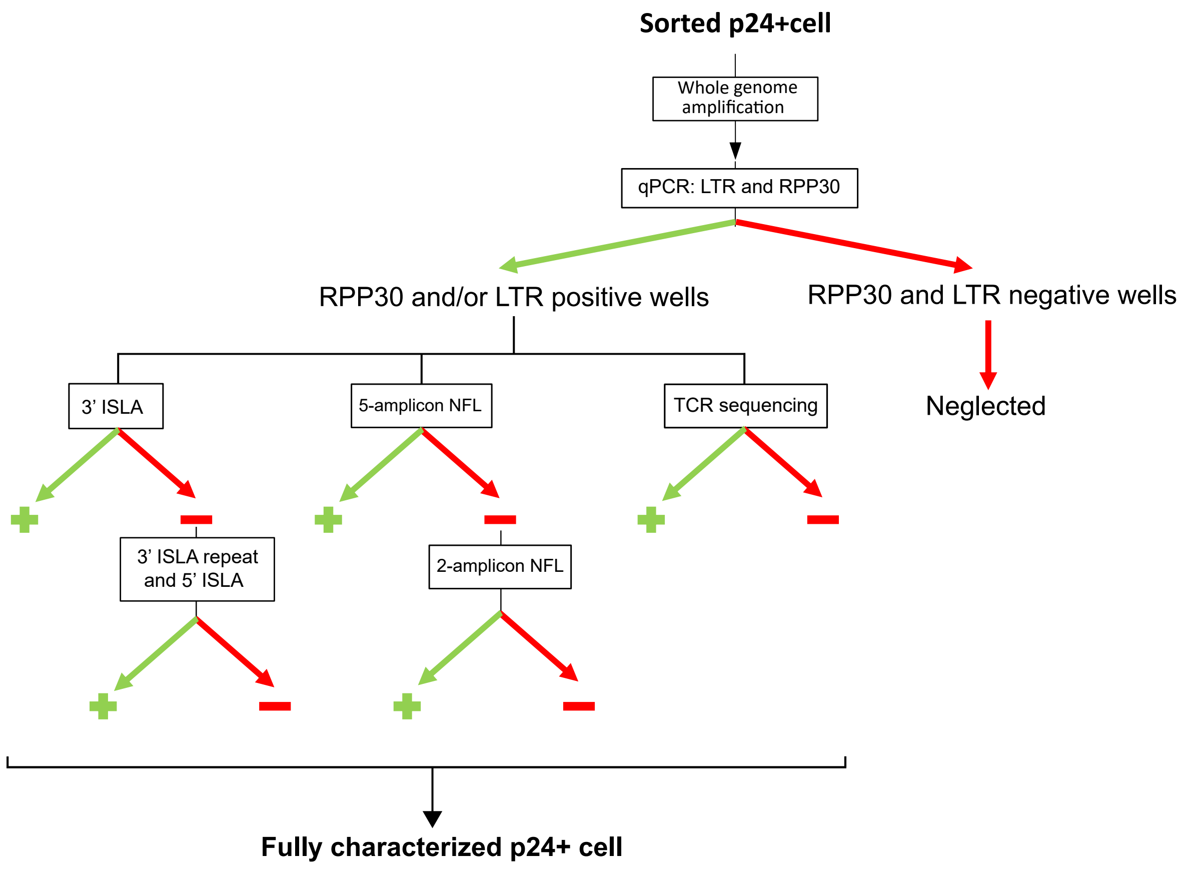

**Supplementary Fig. 3**

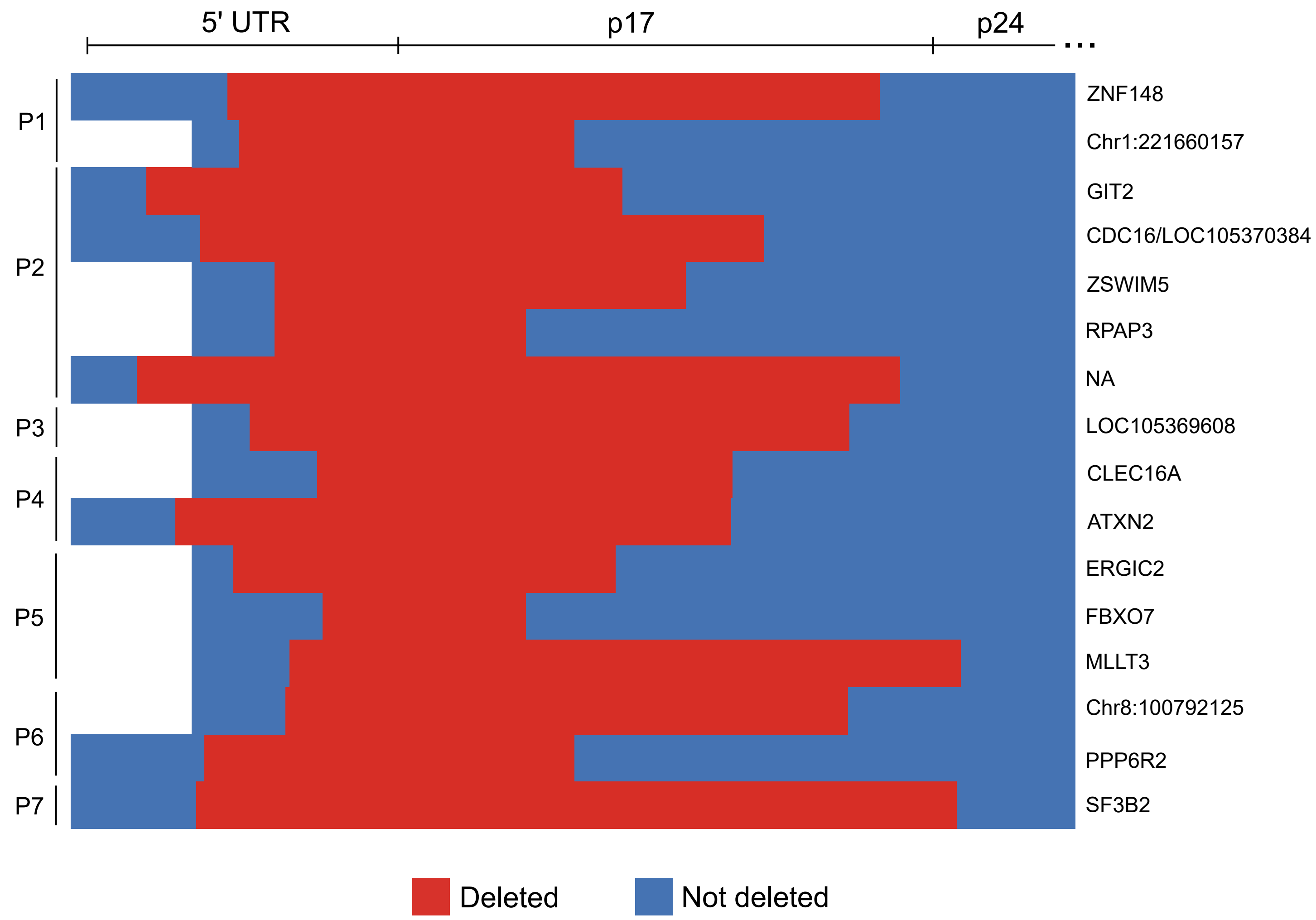

#### Supplementary Fig. 4

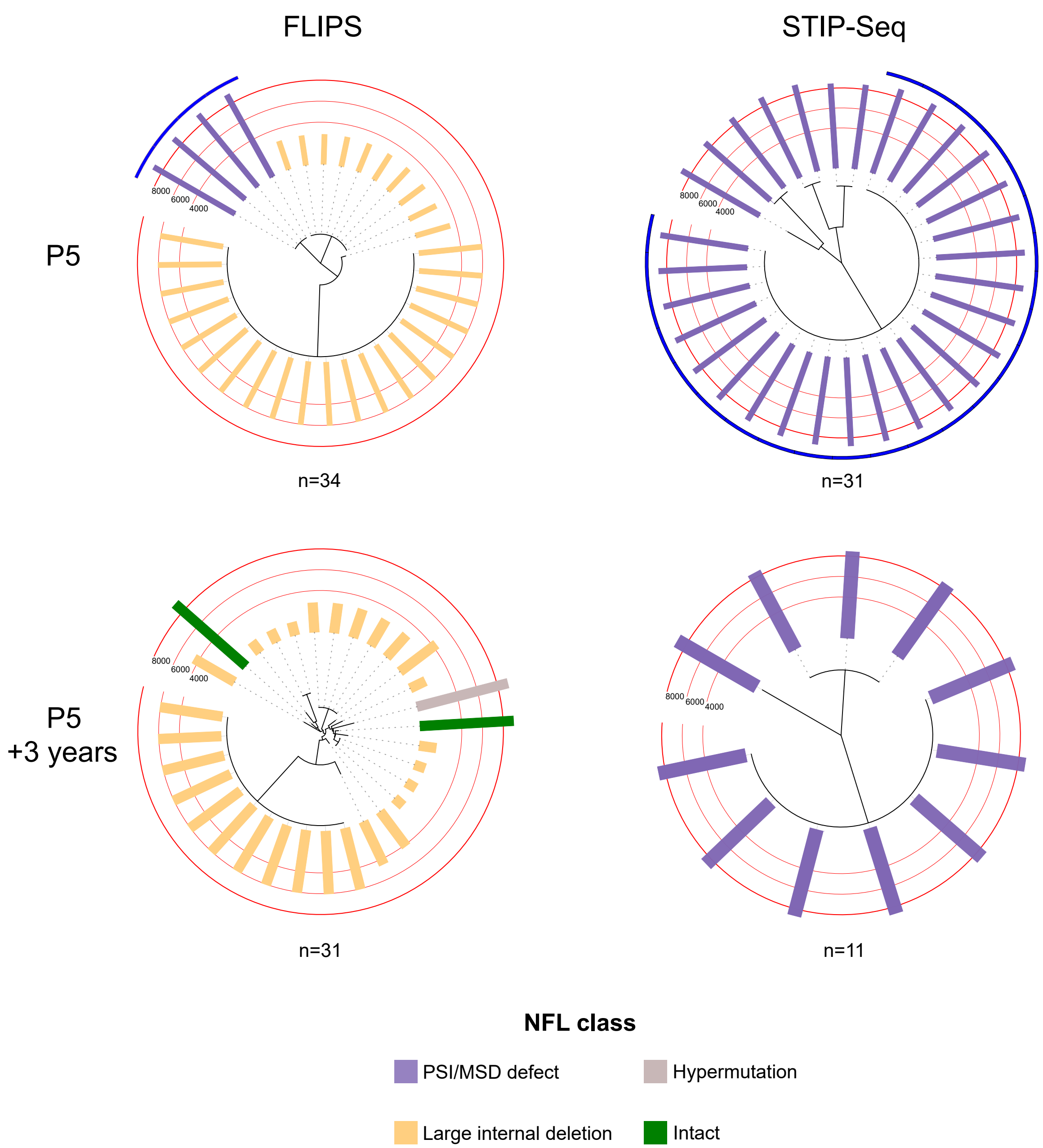

**Supplementary Fig. 5**

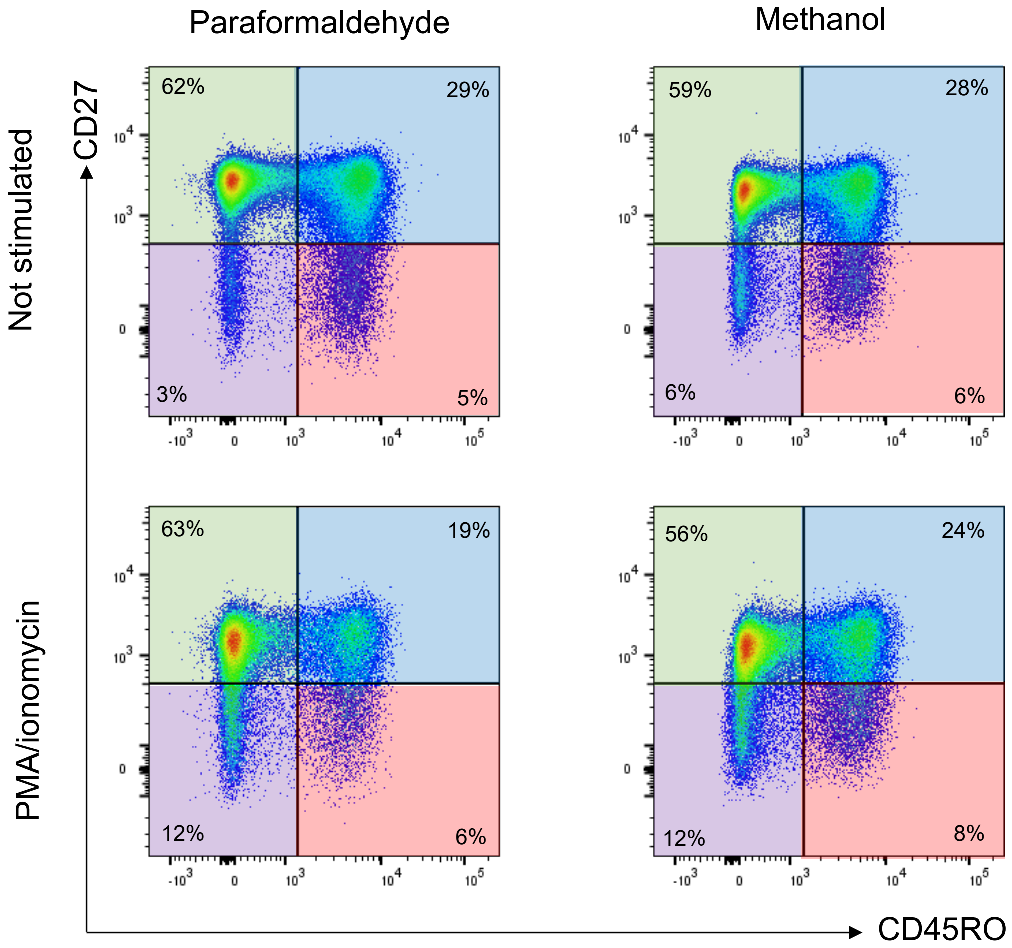

#### Supplementary Fig. 6

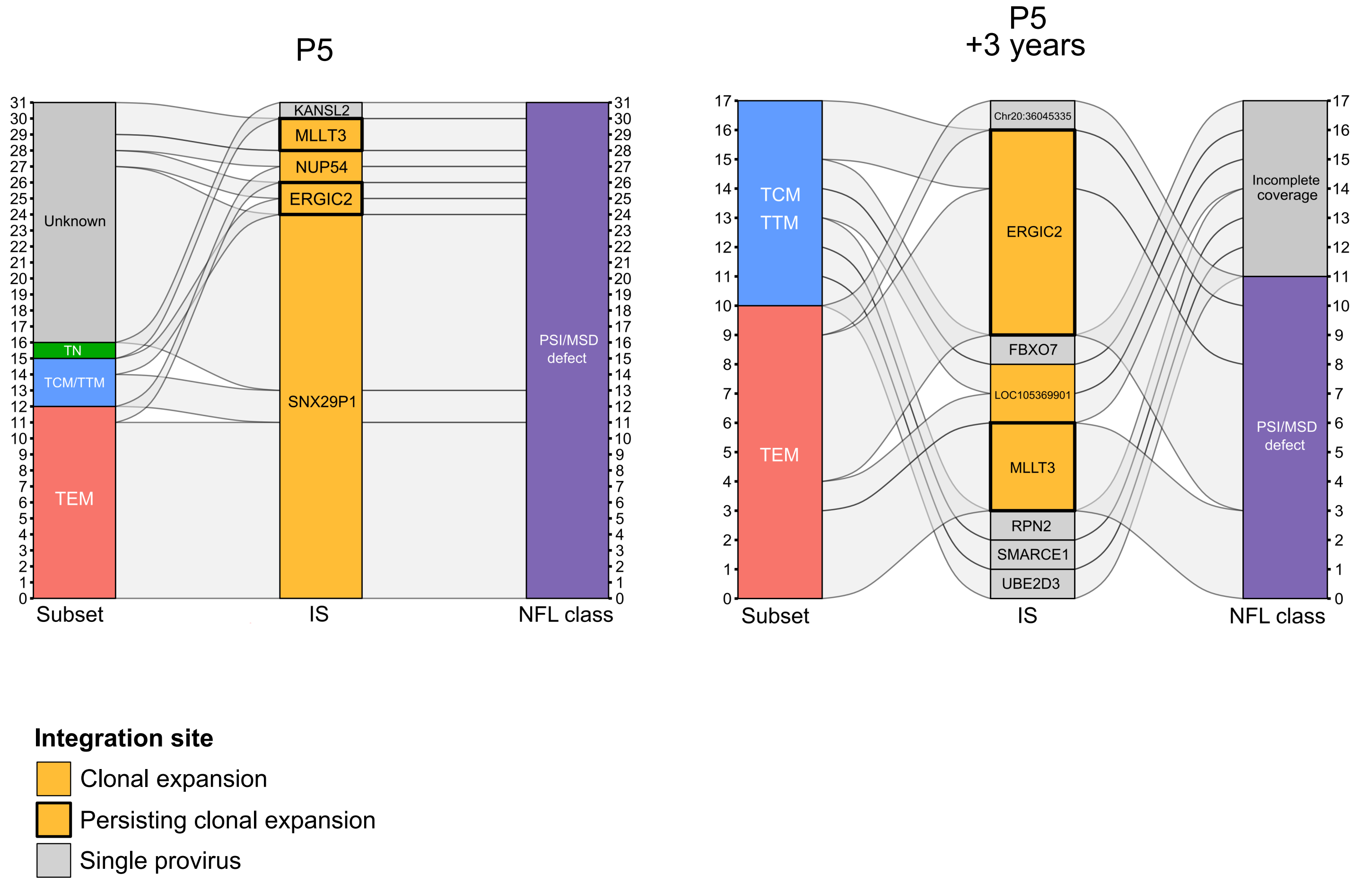

**Supplementary Fig. 7**

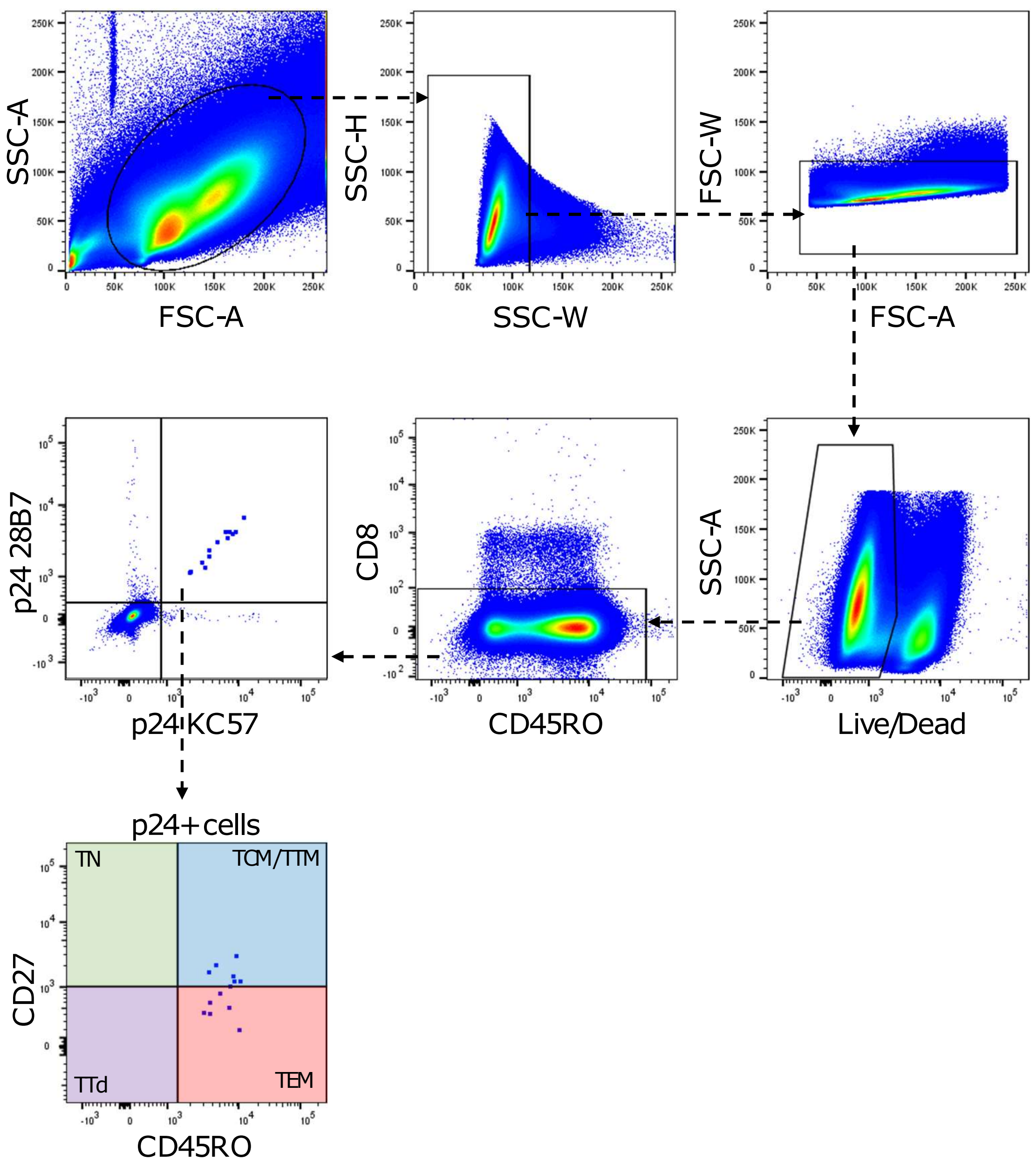

### Supplementary Table 1

| Participant ID | Timepoint | On/off ART | Age | Gender | Year of diagnosis | Year of ART initiation | Time to ART initiation (years) | ART duration (years) | CD4 count (cells/µL) | CD4/CD8 ratio | VL at sampling (copies/ml plasma) | Median p24+ frequency (p24+ cells/million CD4 T cells) |
|---|---|---|---|---|---|---|---|---|---|---|---|---|
| P1 | x | ON | 66 | Male | 1994 | 2001 | 6.5 | 16 | 601 | 0.7 | <40 | 3.5 |
| P2 | x | ON | 61 | Male | 1991 | 1996 | 4.8 | 22 | 1076 | 1.9 | <40 | 9.5 |
| P3 | x | ON | 39 | Male | 2005 | 2012 | 7.8 | 7 | 445 | 2.0 | <40 | 4.2 |
| P4 | x | ON | 57 | Male | 1996 | 1997 | 0.6 | 20 | 911 | 1.0 | < 20 | 2.9 |
| P5 | First | ON | 52 | Male | 2002 | 2002 | 0.4 | 14 | 293 | 0.3 | < 20 | 3.8 |
| P5 | + 3 years | ON | 55 | Male | 2002 | 2002 | 0.4 | 17 | 343 | 0.3 | < 20 | 2.6 |
| P6 | T1 | ON | 54 | Male | 1995 | 1998 | 3.0 | 11 | 736 | 0.6 | < 20 | 1.4 |
| P6 | T2 | OFF | 54 | Male | 1995 | 1998 | 3.0 | 11 | 701 | 0.7 | < 20 | 1.8 |
| P7 | T1 | ON | 37 | Male | 2008 | 2015 | 6.9 | 2 | 911 | 0.9 | < 20 | 0.9 |
| P7 | T2 | OFF | 37 | Male | 2008 | 2015 | 6.9 | 2 | 985 | 1.6 | < 20 | 0.8 |
| P8 | T1 | ON | 52 | Male | 2013 | 2013 | 0.4 | 3 | 714 | 1.3 | < 20 | 2.7 |

### Supplementary Table 2

| Participant | Integration site | Number detected | 5-amplicon NFL | | | | | 2-amplicon NFL | | 4-amplicon NFL | | | |
|---|---|---|---|---|---|---|---|---|---|---|---|---|---|
| | | | A1mod2 (638-2724) | pol (2011-3798) | C (3626-5980) | A2 (5549-7760) | B2 (7652-9610) | Left half (581-5783) | Right half (5088-9602) | Frag 1 (634-3500) | Frag 2 (1870-5248) | Frag 3 (4133-7338) | Frag 4 (6445-9632) |
| P1 | ZNF148 | 4 | | | | | | | | | | | |
| P1 | Chr1:221660157 | 1 | | | | | | | | | | | |
| P1 | ADARB1 | 3 | | | | | | | | | | | |
| P1 | Chr9:61817567 | 1 | | | | | | | | | | | |
| P1 | STAT5B | 1 | | | | | | | | | | | |
| P2 | ITGB1 | 10 | | | | | | | | | | | |
| P2 | NA | 1 | | | | | | | | | | | |
| P2 | GALM* | 1 | | | | | | | | | | | |
| P2 | KHDRBS1 | 1 | | | | | | | | | | | |
| P2 | CDC16/LOC105370384* | 1 | | | | | | | | | | | |
| P2 | GIT2 | 1 | | | | | | | | | | | |
| P2 | MGAT4A | 1 | | | | | | | | | | | |
| P2 | RSPRY1 | 1 | | | | | | | | | | | |
| P2 | ZSWIM5 | 1 | | | | | | | | | | | |
| P2 | AP2A2 | 1 | | | | | | | | | | | |
| P2 | NSD1 | 1 | | | | | | | | | | | |
| P2 | RASA3 | 1 | | | | | | | | | | | |
| P2 | CPEB4 | 1 | | | | | | | | | | | |
| P2 | MAN2A1 | 1 | | | | | | | | | | | |
| P2 | RPAP3 | 1 | | | | | | | | | | | |
| P3 | TNRC6B | 16 | | | | | | | | | | | |
| P3 | LOC105369608 | 1 | | | | | | | | | | | |
| P3 | STAT5B | 4 | | | | | | | | | | | |
| P3 | CD200R1 | 3 | | | | | | | | | | | |
| P3 | Chr2:54321973* | 1 | | | | | | | | | | | |
| P3 | THOC5* | 1 | | | | | | | | | | | |
| P4 | CLEC16A | 1 | | | | | | | | | | | |
| P4 | CMAHP | 1 | | | | | | | | | | | |
| P4 | FCGRT | 1 | | | | | | | | | | | |
| P4 | Chr17:8974901 | 8 | | | | | | | | | | | |
| P4 | ZNF274 | 1 | | | | | | | | | | | |
| P4 | DNAH6 | 1 | | | | | | | | | | | |
| P4 | CD27-AS1 | 1 | | | | | | | | | | | |
| P4 | PLCG1 | 1 | | | | | | | | | | | |
| P4 | CPNE1 | 1 | | | | | | | | | | | |
| P4 | USP12/RBBP8P2 | 2 | | | | | | | | | | | |
| P4 | NUTM2F/HIATL1 | 2 | | | | | | | | | | | |
| P4 | Chr9:136929821 | 1 | | | | | | | | | | | |
| P4 | ATXN2 | 1 | | | | | | | | | | | |
| P5 | SNX29P1/P2 | 24 | | | | | | | | | | | |
| P5 | MLLT3 | 5 | | | | | | | | | | | |
| P5 | ERGIC2 | 8 | | | | | | | | | | | |
| P5 | KANSL2 | 1 | | | | | | | | | | | |
| P5 | NUP54 | 2 | | | | | | | | | | | |
| P5 | UBE2D3 | 1 | | | | | | | | | | | |
| P5 | FBXO7 | 1 | | | | | | | | | | | |
| P5 | SMARCE1 | 1 | | | | | | | | | | | |
| P5 | RPN2 | 1 | | | | | | | | | | | |
| P5 | LOC105369901 | 2 | | | | | | | | | | | |
| P5 | Chr20:36045335 | 1 | | | | | | | | | | | |
| P6 | PPP6R2 | 1 | | | | | | | | | | | |
| P6 | STAT5B | 12 | | | | | | | | | | | |
| P6 | Chr8:100792125 | 4 | | | | | | | | | | | |
| P6 | VMP1 | 3 | | | | | | | | | | | |
| P6 | CIT | 1 | | | | | | | | | | | |
| P7 | Chr17:77978920 | 3 | | | | | | | | | | | |
| P7 | Chr17:7545670 | 5 | | | | | | | | | | | |
| P7 | KCNA3 | 7 | | | | | | | | | | | |
| P7 | SF3B2 | 1 | | | | | | | | | | | |
| P7 | NA | 1 | | | | | | | | | | | |
| P7 | VMP1 | 2 | | | | | | | | | | | |
| P8 | SMG1P2 | 6 | | | | | | | | | | | |

Amplicon name (HXB2 coordinates)

NA = not available
*Sequencing failed

#### Supplementary Table 3

| Participant | Chrom. | Position | Strand | Gene | Orientation | Number detected |
|---|---|---|---|---|---|---|
| P1 | 3 | 125291341 | - | ZNF148 | Same | 4 |
| P1 | 1 | 221660157 | - | NA | NA | 1 |
| P1 | 21 | 45087397 | + | ADARB1 | Opposite | 3 |
| P1 | 9 | 61817567 | - | NA | NA | 1 |
| P1 | 17 | 42253537 | + | STAT5B* | Opposite | 1 |
| P2 | 10 | 32919286 | + | ITGB1 | Opposite | 10 |
| P2 | 2 | 38669512 | - | GALM | Opposite | 1 |
| P2 | 1 | 32030557 | - | KHDRBS1 | Opposite | 1 |
| P2 | 13 | 114258553 | - | CDC16/LOC105370384 | Opposite/Same | 1 |
| P2 | 12 | 109948596 | + | GIT2 | Opposite | 1 |
| P2 | 2 | 98720877 | + | MGAT4A | Opposite | 1 |
| P2 | 16 | 57192144 | + | RSPRY1 | Same | 1 |
| P2 | 1 | 45187297 | + | ZSWIM5 | Opposite | 1 |
| P2 | 11 | 941911 | - | AP2A2 | Opposite | 1 |
| P2 | 5 | 177176435 | + | NSD1* | Same | 1 |
| P2 | 13 | 114093356 | + | RASA3* | Opposite | 1 |
| P2 | 5 | 173905709 | - | CPEB4* | Opposite | 1 |
| P2 | 5 | 109851902 | + | MAN2A1 | Same | 1 |
| P2 | 12 | 47677422 | + | RPAP3 | Opposite | 1 |
| P3 | 22 | 40309709 | + | TNRC6B | Same | 16 |
| P3 | 12 | 3952756 | - | LOC105369608 | Same | 1 |
| P3 | 17 | 42258445 | + | STAT5B* | Opposite | 4 |
| P3 | 3 | 112951052 | + | CD200R1 | Opposite | 3 |
| P3 | 2 | 54321973 | - | NA | NA | 1 |
| P3 | 22 | 29509957 | + | THOC5* | Opposite | 1 |
| P4 | 16 | 11119179 | - | CLEC16A | Opposite | 1 |
| P4 | 6 | 25130744 | + | CMAHP | Opposite | 1 |
| P4 | 19 | 49517975 | - | FCGRT | Opposite | 1 |
| P4 | 17 | 8974901 | + | NA | NA | 8 |
| P4 | 19 | 58195268 | - | ZNF274 | Opposite | 1 |
| P4 | 2 | 84652442 | - | DNAH6 | Opposite | 1 |
| P4 | 12 | 6439170 | + | CD27-AS1 | Opposite | 1 |
| P4 | 20 | 41167601 | - | PLCG1* | Opposite | 1 |
| P4 | 20 | 35637188 | + | CPNE1 | Opposite | 1 |
| P4 | 13 | 27077455 | - | USP12/RBBP8P2 | Same/Opposite | 2 |
| P4 | 9 | 94413057 | + | NUTM2F/HIATL1 | Same/Opposite | 2 |
| P4 | 9 | 136929821 | - | NA | NA | 1 |
| P4 | 12 | 111484919 | - | ATXN2 | Same | 1 |
| P5 | 16 | 21383557 | - | SNX29P1/P2 | Opposite | 24 |
| P5 | 9 | 20608766 | + | MLLT3* | Opposite | 5 |
| P5 | 12 | 29347700 | + | ERGIC2 | Opposite | 8 |
| P5 | 12 | 48680541 | + | KANSL2 | Opposite | 1 |
| P5 | 4 | 76124580 | - | NUP54 | Same | 2 |
| P5 | 4 | 102812614 | + | UBE2D3 | Opposite | 1 |
| P5 | 22 | 32494019 | - | FBXO7 | Opposite | 1 |
| P5 | 17 | 40641871 | + | SMARCE1* | Opposite | 1 |
| P5 | 20 | 37227689 | + | RPN2 | Same | 1 |
| P5 | 12 | 91925519 | + | LOC105369901 | Opposite | 2 |
| P5 | 20 | 36045335 | + | NA | NA | 1 |
| P6 | 22 | 50358570 | - | PPP6R2 | Opposite | 1 |
| P6 | 17 | 42264648 | + | STAT5B* | Opposite | 12 |
| P6 | 8 | 100792125 | + | NA | NA | 4 |
| P6 | 17 | 59803733 | - | VMP1*[†] | Opposite | 3 |
| P6 | 12 | 119761801 | - | CIT[†] | Same | 1 |
| P7 | 17 | 77978920 | - | NA | NA | 3 |
| P7 | 17 | 7545670 | - | NA | NA | 5 |
| P7 | 1 | 110655131 | - | KCNA3 | Same | 7 |
| P7 | 11 | 66064686 | - | SF3B2 | Opposite | 1 |
| P7 | 17 | 59803547 | - | VMP1* | Opposite | 2 |
| P8 | 16 | 29558266 | - | SMG1P2 | Same | 6 |

[†]Only retrieved at T2, during analytical treatment interruption

NA = not applicable

#### Supplementary Table 4

| Participant | Integration site | Number detected | TRBV | CDR3 (AA) | TRBJ | Specificity |
|---|---|---|---|---|---|---|
| P1 | ZNF148 | 4 | TRBV7-3 | GRNQPQHF | TRBJ1-5 | NA |
| P1 | Chr1:221660157 | 1 | TRBV18 | CASSPGGLAETQFF | TRBJ2-6 | NA |
| P1 | ADARB1 | 3 | TRBV27 | CASSFFLNNHPPHF | TRBJ1-5 | NA |
| P1 | Chr9:61817567 | 1 | TRBV7-9 | CASSQGDGYGYTF | TRBJ1-2 | *M. tuberculosis* |
| P1 | STAT5B | 1 | TRBV9 | CASSVVNRYAETQFF | TRBJ2-5 | NA |
| P2 | ITGB1 | 9 | TRBV20-1 | CSAGPSNQPQHF | TRBJ1-5 | NA |
| P2 | NA | 1 | NA | NA | NA | NA |
| P2 | GALM | 1 | TRBV3-1 | CASSQVTSGGARETHFF | TRBJ2-5 | NA |
| P2 | KHDRBS1 | 1 | TRBV5-5 | CASSSRTGGNEQFF | TRBJ2-1 | NA |
| P2 | CDC16/LOC105370384 | 1 | NA | NA | NA | NA |
| P2 | GIT2 | 1 | TRBV5-4 | CASSLQGASYNEQFF | TRBJ2-1 | NA |
| P2 | MGAT4A | 1 | TRBV20-1 | CSARVRTSGYEQYF | TRBJ2-7 | NA |
| P2 | RSPRY1 | 1 | TRBV5-4 | CASSLVGAPPLNTEAFF | TRBJ1-1 | NA |
| P2 | ZSWIM5 | 1 | TRBV29-1 | CSVLPGQTEAFF | TRBJ1-1 | NA |
| P2 | AP2A2 | 1 | NA | NA | NA | NA |
| P2 | NSD1 | 1 | TRBV30 | CAWSARTDTQYF | TRBJ2-3 | NA |
| P2 | RASA3 | 1 | NA | NA | NA | NA |
| P2 | CPEB4 | 1 | NA | NA | NA | NA |
| P2 | MAN2A1 | 1 | NA | NA | NA | NA |
| P2 | RPAP3 | 1 | TRBV14 | CASSKGLAGGVTTDTQYF | TRBJ2-3 | NA |
| P3 | TNRC6B | 16 | TRBV5-1 | CASSLEAGGNHPLF | TRBJ1-3 | NA |
| P3 | LOC105369608 | 1 | NA | NA | NA | NA |
| P3 | STAT5B | 4 | TRBV4-2 | CASSQDEGYGYTF | TRBJ1-2 | Influenza |
| P3 | CD200R1 | 3 | TRBV9 | CASSPQGLNTEAFF | TRBJ1-1 | CMV, influenza, *M. tuberculosis* |
| P3 | Chr2:54321973 | 1 | TRBV15 | CATSRAQSNEKLFF | TRBJ1-4 | NA |
| P3 | THOC5 | 1 | NA | NA | NA | NA |
| P4 | CLEC16A | 1 | NA | NA | NA | NA |
| P4 | CMAHP | 1 | TRBV7-9 | CASSPGTYNSPLHF | TRBJ1-6 | *M. tuberculosis* |
| P4 | FCGRT | 1 | TRBV20-1 | CSAQGPQGPQHF | TRBJ1-5 | NA |
| P4 | Chr17:8974901 | 8 | TRBV20-1 | CSARVRDRPYEQYF | TRBJ2-7 | *M. tuberculosis* |
| P4 | ZNF274 | 1 | TRBV15 | CATGTLAGRTLNTEAFF | TRBJ1-1 | NA |
| P4 | DNAH6 | 1 | NA | NA | NA | NA |
| P4 | CD27-AS1 | 1 | NA | NA | NA | NA |
| P4 | PLCG1 | 1 | TRBV7-9 | CASSQGSGEGETQFF | TRBJ2-5 | NA |
| P4 | CPNE1 | 1 | NA | NA | NA | NA |
| P4 | USP12/RBBP8P2 | 2 | TRBV11-2 | CASRRNAGTSDEQYF | TRBJ2-7 | NA |
| P4 | NUTM2F/HIATL1 | 2 | TRBV19 | CASSIGQGSNEELFF | TRBJ1-4 | NA |
| P4 | Chr9:136929821 | 1 | TRBV6-1 | CASSLKPRGANYGYYF | TRBJ1-2 | NA |
| P4 | ATXN2 | 1 | NA | NA | NA | NA |
| P5 | SNX29P1/P2 | 24 | TRBV7-9 | CASSRYRGR#TEAFF | TRBJ1-1 | NA |
| P5 | MLLT3 | 5 | NA | NA | NA | NA |
| P5 | ERGIC2 | 8 | TRBV11-2 | CASSLDGDSPLPF | TRBJ1-6 | NA |
| P5 | KANSL2 | 1 | TRBV18 | CASSPEVGSIGEQYF | TRBJ2-7 | NA |
| P5 | NUP54 | 2 | TRBV20-1 | CSADLGHRYPLF | TRBJ2-3 | NA |
| P5 | UBE2D3 | 1 | NA | NA | NA | NA |
| P5 | FBXO7 | 1 | NA | NA | NA | NA |
| P5 | SMARCE1 | 1 | TRBV5-1 | CASSSPGQGYEQYF | TRBJ2-7 | NA |
| P5 | RPN2 | 1 | TRBV6-5 | CASRRKGGRWDTTAFF | TRBJ1-1 | NA |
| P5 | LOC105369901 | 2 | NA | NA | NA | NA |
| P5 | Chr20:36045335 | 1 | TRBV6-6 | CAASSGGVLPPHFF | TRBJ1-6 | NA |
| P6 | PPP6R2 | 1 | TRBV6-1 | CASSEGWDDQQPPHFF | TRBJ1-4 | NA |
| P6 | STAT5B | 12 | TRBV19 | CASSIAGRAFNASTF | TRBJ1-2 | NA |
| P6 | Chr8:100792125 | 4 | TRBV2 | CASSPNRGRGYTF | TRBJ1-2 | CMV |
| P6 | VMP1 | 3 | NA | NA | NA | NA |
| P6 | CIT | 1 | NA | NA | NA | NA |
| P7 | Chr17:77978920 | 3 | TRBV28 | CATSFCGGRAADVLYF | TRBJ2-4 | NA |
| P7 | Chr17:7545670 | 5 | TRBV5-5 | CASSLGAGTGGTYGYTF | TRBJ1-2 | NA |
| P7 | KCNA3 | 7 | TRBV7-2 | CASSLKTGGYEQYF | TRBJ2-7 | *M. tuberculosis* |
| P7 | SF3B2 | 1 | TRBV12-3 | CASSLRDAAVAFF | TRBJ1-1 | NA |
| P7 | NA | 1 | TRBV5-4 | CASSSKRGSTDTQYF | TRBJ2-3 | *M. tuberculosis* |
| P7 | VMP1 | 2 | TRBV28 | CASRPRRTYCYYF | TRBJ2-7 | NA |
| P8 | SMG1P2 | 6 | TRBV18 | CASSPEVGSIGEQYF | TRBJ2-7 | NA |

NA = not available

#### Supplementary Table 5

| Assay | Amplicon | Round | HXB2 coordinates | Forward/reverse | Primer | Sequence (5' to 3') |
|---|---|---|---|---|---|---|
| 5-amplicon NFL | A1mod2 | 1 | 623-3333 | Forward | U5-623F | AAATCTCTAGCAGTGGCGCCCGAACAG |
| | | | | Reverse | NE1 | CCACTAACTTCTGTATGTCATTGACAGTCCAGCT |
| | | 2 | 638-2724 | Forward | U5-638F | GCGCCCGAACAGGGACYTGAAARCGAAAG |
| | | | | Reverse | ProC- | GAGTATTGTATGGATTTTCAGGCCCAAT |
| | pol | 1 | 1981-3859 | Forward | 5CP1 | GAAGGGCACACAGCCAGAAATTGCAGGG |
| | | | | Reverse | RT3.1 | GCTCCTACTATGGGTTCTTTCTCTAACTGG |
| | | 2 | 2011-3798 | Forward | 2.5 | CCTAGGAAAAAGGGCTGTTGGAAATGTGG |
| | | | | Reverse | RT3798R | CAAACTCCCACTCAGGAATCCA |
| | C | 1 | 3597-6004 | Forward | RT3597mixF | AAAACAGGAAARTATGCAA |
| | | | | Reverse | SC05R | AGCTCTTCGTCGCTGTCTCCGCTT |
| | | 2 | 3626-5980 | Forward | RT3626F | TGCCCACACTAATGATGTAA |
| | | | | Reverse | SC02R | CTTCCTGCCATAGGAGATGCCTA |
| | A2 | 1 | 5450-7817 | Forward | VP5450F | CAGGACATAACAAGGTAGGATC |
| | | | | Reverse | CO602 | GCCCATAGTGCTTCCTGCTGCTCCCAAGAACC |
| | | 2 | 5549-7760 | Forward | VP5549F | AGAGGATAGATGGAACAAGCCCCAG |
| | | | | Reverse | V3CR | TGCTCTTTTTTCTCTCTSCACCACT |
| | B2 | 1 | 7626-9628 | Forward | GP41Fo | TTCAGACCTGGAGGAGGAGATAT |
| | | | | Reverse | 3LTRi | TCAAGGCAAGCTTTATTGAGGCTTAA |
| | | 2 | 7652-9610 | Forward | GP41Fi | GGACAATTGGAGAAGTGAATTAT |
| | | | | Reverse | 3UTRi | AGGCTTAAGCAGTGGGTTCCCTAG |
| 2-amplicon NFL | Left half | 1 | 544-5968 | Forward | F544 | TTAAGCCTCAATAAAGCTTGCCTTGAG |
| | | | | Reverse | R5968 | TGTCTYCKCTTCTTCCTGCCATAG |
| | | 2 | 581-5783 | Forward | F581 | GTGTGCCCGTCTGTTGTGTGACTC |
| | | | | Reverse | R5783 | AATGCCTATTCTGCTATGTYGACACC |
| | Right half | 1 | 5066-9665 | Forward | F5066alt1 | TATGGAAAACAGATGGCAGGTGMTGRT |
| | | | | Reverse | R9665 | GTCTGAGGGATCTCTAGWTACCAGA |
| | | 2 | 5088-9602 | Forward | F5088alt1 | GATTGTGTGGCARGTAGACAGRATG |
| | | | | Reverse | R9602 | CAAGGCAAGCTTTATTGAGGCTTAAS |
| 4-amplicon NFL | NFL | 1 | 611-9675 | Forward | 611(+) | AGTCAGTGTGGAAAATCTCT*A*G |
| | | | | Reverse | 9675(-) | GAGGGATCTCTAGTTACCAG*A*G |
| | Frag 1 | 2 | 634-3500 | Forward | 634(+) | AGTGGCGCCCGAACAGGGAC |
| | | | | Reverse | 3500(-) | CTATTAAGTATTTTGATGGGTCATAA |
| | Frag 2 | 2 | 1870-5248 | Forward | 1870(+) | GAGTTTTGGCTGAGGCAATGAG |
| | | | | Reverse | 5248(-) | TCTCCTGTATGCAGACCCCA |
| | Frag 3 | 2 | 4133-7338 | Forward | 4133(+) | GGAAAAGGTCTACCTGGCATG |
| | | | | Reverse | E125(-) | CAATTTCTGGGTCCCCTCCTGAGG |
| | Frag 4 | 2 | 6445-9632 | Forward | E30(+) | GTGTACCCACAGACCCCAGCCCACAAG |
| | | | | Reverse | R-519(-) | GCACTCAAGGCAAGCTTTATTGAGGCTTA |
